# Outer membrane remodeling via lipid-peptidoglycan crosstalk enables lipooligosaccharide-deficient colistin resistance

**DOI:** 10.64898/2026.05.04.722707

**Authors:** Roberto Jhonatan Olea-Ozuna, Berenice Furlan, Hanling Gong, Orietta Massidda, Joseph M. Boll

## Abstract

Gram-negative bacteria rely on an asymmetric outer membrane (OM) for barrier integrity, with phospholipids confined to the inner leaflet and glycolipids such as lipopolysaccharide (LPS) or lipooligosaccharide (LOS) forming the outer leaflet. Although LPS/LOS was long considered essential, recent findings challenge this view, leaving the mechanistic basis and evolutionary flexibility unclear. Here, we identify lipid asymmetry as a structural checkpoint that governs access to LOS-independent survival. Using *Acinetobacter baumannii* as a model, we show that disrupting retrograde phospholipid transport and surface phospholipid degradation destabilizes OM lipid balance, creating a permissive state that enables emergence of LOS-deficient, colistin-resistant variants. Integrated lipidomic and transcriptomic analyses reveal a staged remodeling program that reinforces lipoprotein scaffolds, rewires peptidoglycan synthesis, and expands trafficking pathways to stabilize a glycolipid-free envelope. Critically, loss of LOS coincides with sharp repression of PBP1A, and maintaining its activity blocks adaptation, demonstrating interdependence between OM and peptidoglycan homeostasis. We propose a three-state model—basal, permissive, adapted—that explains how envelope architecture gates evolutionary trajectories to antibiotic resistance.

**Significance:** Colistin is a last-resort antibiotic targeting lipopolysaccharide (LPS) or lipooligosaccharide (LOS) in Gram-negative pathogens, and many emerging antimicrobials aim to inhibit LPS/LOS biosynthesis and transport. Resistance usually arises via lipid A modification, which preserves LPS/LOS while reducing colistin binding. Resistance to colistin can also arise via complete loss of LOS, which occurs in some *Acinetobacter baumannii* strains but is constrained in others, such as strain ATCC 17978. Here, we demonstrate that LOS essentiality is not fixed but dictated by outer membrane architecture. Disrupting phospholipid homeostasis creates a permissive envelope that allows LOS-deficient, colistin-resistant variants to emerge, while reducing peptidoglycan synthesis further promotes this state. These findings identify lipid asymmetry as a structural checkpoint in resistance evolution and suggest that preserving envelope homeostasis could limit bacterial escape from colistin and guide strategies for next-generation antibiotic development.

## Introduction

The Gram-negative outer membrane (OM) is an asymmetric bilayer that underpins barrier integrity and bacterial survival (1, 2). Phospholipids (PLs) are normally confined to the inner leaflet, whereas lipopolysaccharide (LPS) or lipooligosaccharide (LOS) glycolipids form the outer leaflet that confers low permeability and a characteristic anionic surface charge (3, 4). This lipid asymmetry is actively maintained by coordinated transport and ativey-control systems. Defects in these systems increase OM permeability and susceptibility to environmental and antibiotic stress (5–7). A long-standing assumption has been that LPS/LOS is essential for viability in Gram-negative species (8), yet multiple lineages can survive with severely reduced or absent LPS/LOS under defined conditions (9–12). These observations raise a general question: what structural principles of the envelope gate access to glycolipid-independent survival, and how do cells functionally traverse that gate under antibiotic selection?

Here, we identify lipid asymmetry as a structural checkpoint that governs entry into glycolipid-independent states and reveal how peptidoglycan (PG) synthesis sets the mechanical tolerance for this transition. Using *Acinetobacter baumannii* as a tractable model, we show that simultaneous disruption of retrograde PL transport (Mla) and surface PL degradation (PldA) destabilizes OM lipid balance to create a permissive but fragile envelope in which LOS loss becomes accessible under colistin selection. This trajectory bypasses the canonical route of lipid A modification and instead depends on the mechanical and compositional state of the envelope. Integrated lipidomics and transcriptomics reveal an ordered remodeling program that (i) reinforces protein and lipoprotein scaffolds, (ii) rewires PG synthesis, and (iii) expands OM trafficking to stabilize a glycolipid-free surface. Mechanistically, we find that PBP1A (encoded by *mrcA*), a bifunctional PG synthase, is sharply repressed upon LOS loss, and that maintaining PBP1A activity blocks emergence of LOS-deficient, colistin-resistant variants—demonstrating interdependence between OM asymmetry and PG biosynthesis in defining envelope viability (12–15).

From these data, we derive a three-state conceptual model (**Figure 1**) that generalizes across Gram-negative envelopes. In the (1) basal state (LOS-dependent), lipid asymmetry is actively maintained enforcing glycolipid essentiality and ensuring low OM permeability, with colistin targeting lipid A. The (2) permissive state (asymmetry collapsed) arises when PL homeostasis (Mla/PldA) is disrupted, producing a metastable OM that remains LOS-positive but is structurally weakened, thereby reducing the barrier to glycolipid loss. Finally, in the (3) adapted state (LOS-deficient), cells rebuild the OM around protein/lipoprotein scaffolds, attenuate PG expansion through reduced PBP1A levels, and expand trafficking and stress response pathways, enabling colistin-resistant growth with collateral vulnerabilities.

**Figure 1.**
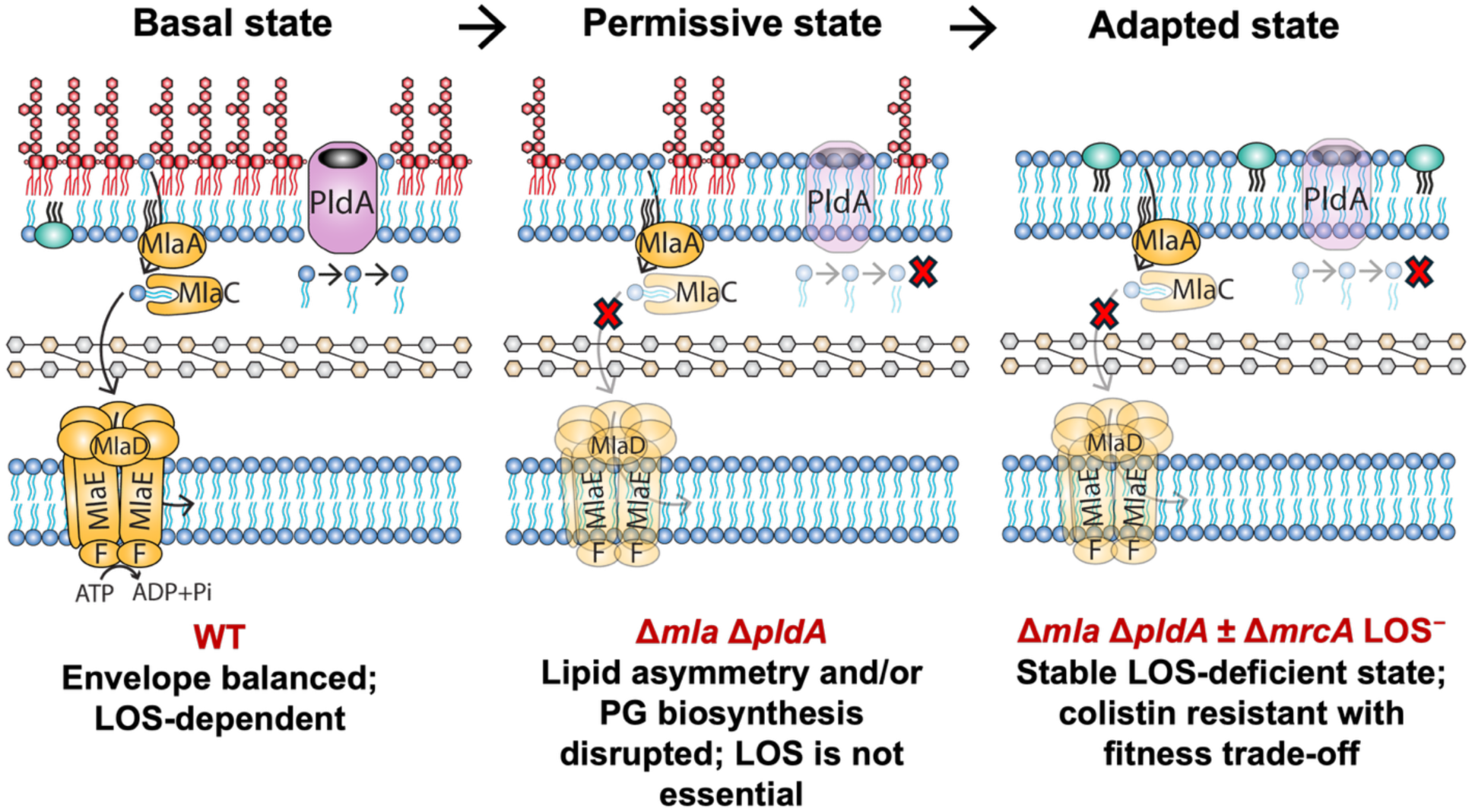
A three-state model of OM remodeling enables LOS-independent colistin resistance in *A. baumannii*. Schematic representation of sequential envelope transitions that facilitate the emergence of colistin-resistant, LOS-deficient variants. <u>Basal state:</u> In wild-type (WT) OM lipid asymmetry is preserved, maintaining low permeability to hydrophilic antibiotics, and LOS remains essential for viability. <u>Permissive state:</u> In the Δ*mla* Δ*pldA* double mutant, loss of PL retrograde transport destabilizes OM lipid asymmetry, generating a metastable envelope that permits LOS loss under colistin selection. <u>Adapted state:</u> LOS-deficient variants arising in this background acquire high-level colistin resistance but display increased susceptibility to antibiotics, altered morphology, and reduced fitness. Further inactivation of *mrcA*, a key PG synthase, enhances envelope destabilization and accelerates transition from the permissive to adapted state. This model illustrates how progressive disruption of lipid and PG homeostasis expands the range of envelope configurations compatible with survival and enables alternative resistance trajectories in *A. baumannii*.

This framework explains when glycolipid loss becomes accessible, how the envelope is stabilized without LOS, and why resistance routes diversify under antibiotic pressure. Importantly, the model articulates general design rules—lipid asymmetry as a gate, PG synthesis as a throttle that modulates how the envelope adapts without LOS—both of which are likely relevant beyond *A. baumannii* to other Gram-negative pathogens exposed to polymyxins or inhibitors of LPS/LOS biogenesis (16–19).

Finally, the work has immediate translational relevance. Colistin directly targets LPS/LOS, and many emerging antimicrobials aim to inhibit LPS/LOS biosynthesis. By revealing the envelope states that enable or block glycolipid-independent survival, our model provides a conceptual foundation for predicting resistance trajectories and for designing adjuvant strategies that preserve lipid asymmetry or sustain PG-driven mechanical constraints—thereby restricting escape from last-resort antibiotics.

## Results

### Disruption of lipid asymmetry reveals vulnerabilities and sensitizes *A. baumannii* to stress

We hypothesized that lipid asymmetry functions as a structural checkpoint enforcing LOS essentiality and that its collapse could create a metastable envelope permissive for LOS loss (**Figure 1**). To test this, we disrupted three systems that contribute to OM integrity and lipid homeostasis. The Mla system and the OM phospholipase PldA are established regulators of PLs homeostasis, mediating retrograde PL transport and outer leaflet PL degradation, respectively (20–25). In contrast, the Tol-Pal complex is mechanistically distinct: it primarily maintains OM-PG connectivity and overall envelope architecture. Although it has been implicated in retrograde PL transport in *Escherichia coli* (26, 27), its role in *A. baumannii* remains less defined (28). Accordingly, while Mla and PldA directly regulate lipid asymmetry, Tol-Pal is more likely to influence OM stability through structural effects on envelope organization. Importantly, disruption of these systems—whether through altered PL trafficking, defective PL turnover, or impaired OM–PG coupling—converges on a common outcome of envelope destabilization and increased surface-exposed PLs (**Figure S1A**).

Each system was disrupted by allelic exchange: the *mlaFEDCB* operon was deleted; *pldA* was removed individually; and the *tol*-*pal* complex, encoded in two adjacent operons (*tolQRA* and *tolB*-*pal*), was deleted as a single functional unit (**Figure S1B**). This genetic framework enabled the construction of all combinations of single, double, and triple mutants for downstream analyses.

To dissect the individual and combined contributions of Mla, PldA, and Tol-Pal to OM homeostasis, we constructed a panel of mutants in *A. baumannii* strain ATCC 17978, including single (Δ*mla*, Δ*pldA*, Δ*tol*-*pal*), double (Δ*mla* Δ*pldA*, Δ*mla* Δ*tol*-*pal*, Δ*pldA* Δ*tol*-*pal*), and triple (Δ*mla* Δ*pldA* Δ*tol*-*pal*) deletions (**Figure 2**). OM stability was assessed by quantifying colony-forming units (CFUs) following exposure to sublethal concentrations of sodium dodecyl sulfate (SDS), which disrupts lipid packing, and ethylenediaminetetraacetic acid (EDTA), which chelates divalent cations required for LOS stabilization (**Figure 2A**).

**Figure 2.**
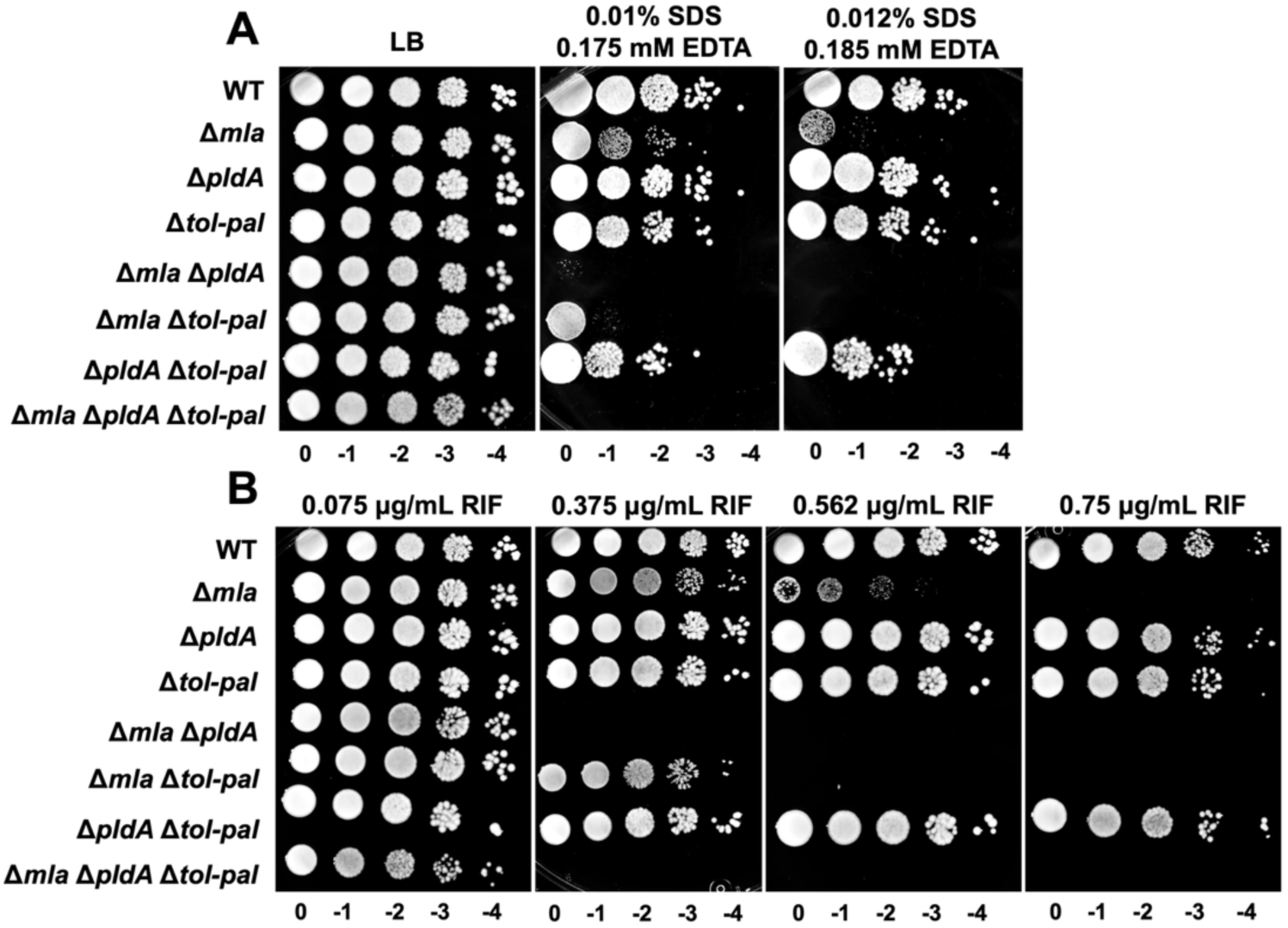
Disruption of OM lipid homeostasis increases sensitivity of *A. baumannii* ATCC 17978 to envelope stress and antibiotic pressure. (A) Viability of wild-type (WT) and mutant strains (including all single, double, and triple deletions) following exposure to SDS and EDTA in nutrient rich LB. (B) Survival of the same strains following exposure to increasing concentrations of rifampicin (RIF). Data shown are representative of three independent experiments.

Under non-stress conditions, all strains grew comparably in rich medium, indicating preserved baseline OM function. However, exposure to mild detergent stress (0.01% SDS, 0.175 mM EDTA) revealed distinct phenotypic differences. Wild-type cells remained fully viable, whereas mutants exhibited graded susceptibility. Single Δ*pldA* or Δ*tol*-*pal* mutants displayed no overt loss of viability and behaved similarly to the wild-type, indicating that disruption of PL degradation or OM tethering alone is insufficient to compromise OM integrity. In contrast, the Δ*mla* mutant exhibited a pronounced loss of viability, underscoring a dominant role for Mla in OM maintenance. Double mutants combining Δ*mla* with either Δ*pldA* or Δ*tol*-*pal* displayed clear synthetic sensitivity, and the triple mutant (Δ*mla* Δp*ldA* Δ*tol*-*pal*) showed severely impaired viability. These effects were exacerbated under higher stress conditions (0.012% SDS, 0.185 mM EDTA), where all Mla-deficient combinations lost viability, consistent with marked OM destabilization.

Plasmid-based expression of *mla* partially restored SDS/EDTA resistance in the Mla-deficient strains, including double and triple mutants (**Figure S2**), confirming that Mla is the primary determinant of OM integrity and cooperates with PldA and Tol-Pal to sustain OM homeostasis and envelope resilience.

To determine whether these structural defects also compromised OM barrier function, we used rifampicin, a hydrophobic antibiotic whose entry is normally restricted by an intact OM, as a probe (**Figure 2B**). At low concentrations (0.075 μg/mL), all strains retained viability. At an intermediate concentration (0.375 μg/mL), mutants lacking both *mla* and *pldA*, as well as the triple mutant, were completely non-viable, while Δ*mla* Δ*tol*-*pal* strain showed marked sensitivity. The Δ*mla* single mutant tolerated rifampicin up to 0.562 μg/mL but became susceptible at 0.75 μg/mL. These results reveal a concentration-dependent hierarchy of OM disruption, with Mla as the primary determinant of barrier function, and PldA and Tol-Pal acting as supportive elements. Complementation with *mla* restored rifampicin resistance in double and triple mutants to intermediate levels (**Figure S2**), further supporting the conclusion that lipid asymmetry is critical for OM impermeability and antibiotic resistance.

Collectively, these findings reveal a multilayered system for OM maintenance in *A. baumannii*, with Mla at the core, supported by the lipid-modifying activity of PldA and the structural role provided by Tol-Pal.

Disruption of this cooperative network sensitizes the OM to detergent and antibiotic stress, exposing vulnerabilities that may be therapeutically exploited to enhance antibiotic efficacy against multidrug-resistant *A. baumannii*.

### Phospholipid remodeling, not LOS depletion, defines the biochemical signature of lipid homeostasis disruption

To assess biochemical consequences of OM lipid disruption, we quantified LOS abundance and total PL composition across wild-type and mutants carrying single, double, and triple deletions of *mla*, *pldA*, and/or *tol*-*pal* (**Figure 3**).

**Figure 3.**
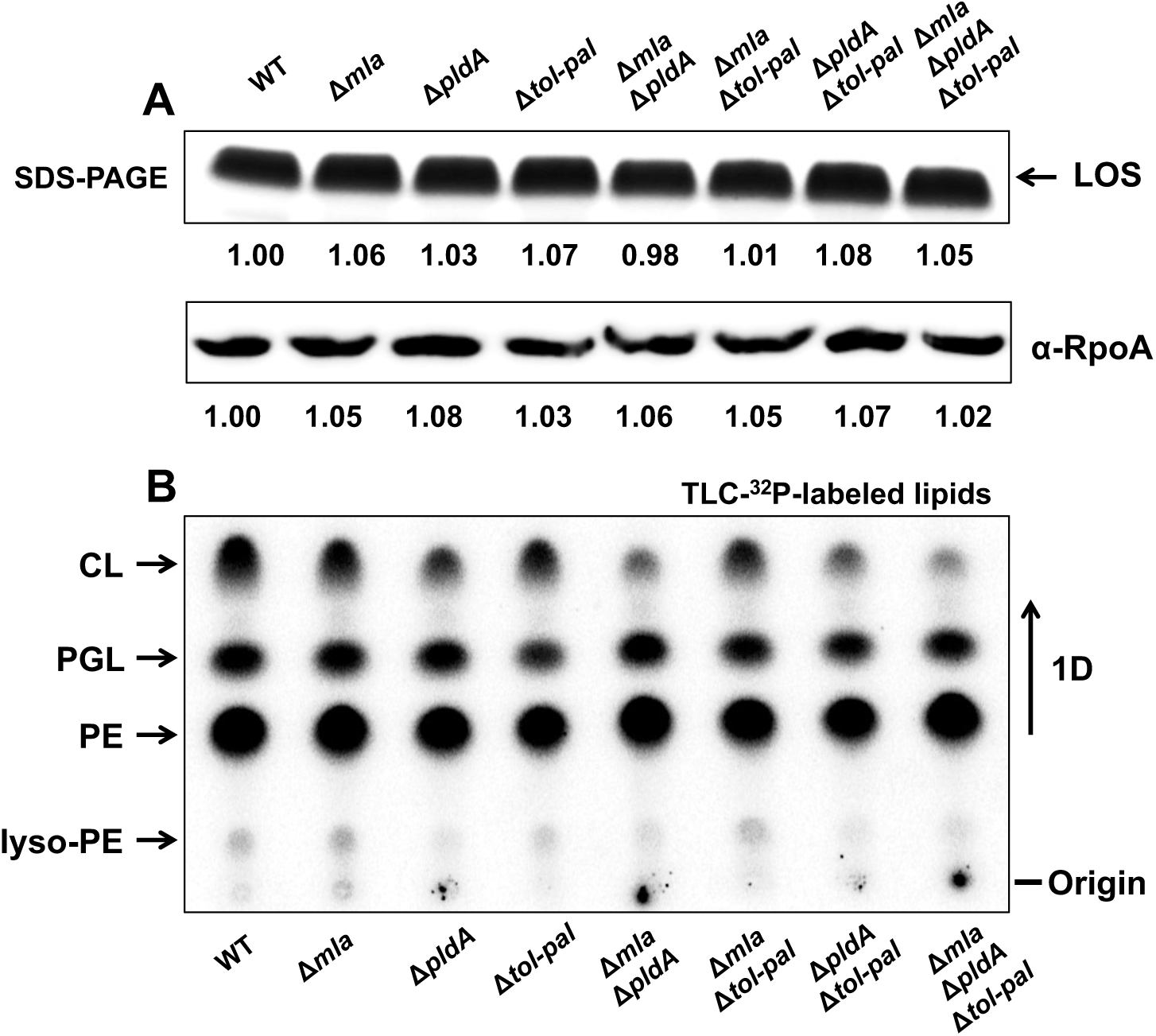
Disruption of OM lipid homeostasis alters PLs composition without affecting LOS abundance. (A) SDS-PAGE analysis of LOS extracted from whole-cell lysates of wild-type (WT) and mutant strains. Samples were normalized to and OD_600_ of 1 prior to extraction. LOS was detected by fluorescent, and band intensities were quantified and normalized to WT (set as 1.0). RpoA (37.62 kDa) was used as a loading control to confirm comparable sample input across strains. (B) PLs labeled with [³²P] were extracted from the same strains and separated by one dimensional (1D) thin-layer chromatography (TLC). The relative abundance of phosphatidylethanolamine (PE), cardiolipin (CL), phosphatidylglycerol (PGL), and lyso-PE is shown as a percentage of total PL signal (see Table 1). Data are representative of three independent experiments.

**Table 1:** Quantification of total PL species by [³²P] labeling and thin-layer chromatography (TLC). Phospholipids were extracted from wild-type (WT) and mutant strains, including single, double, and triple mutants, grown in the presence of [³²P]-orthophosphate and analyzed by TLC. Each value is the mean of three independent experiments ± standard deviation

| Phospholipids | WT | $\Delta mla$ | $\Delta pldA$ | $\Delta tol-pal$ | $\Delta mla \Delta pldA$ | $\Delta mla \Delta tol-pal$ | $\Delta pldA \Delta tol-pal$ | $\Delta mla \Delta pldA \Delta tol-pal$ |
| --- | --- | --- | --- | --- | --- | --- | --- | --- |
| CL | 30.45±0.16 | 27.28±0.10 | 22.31±0.75 | 28.50±0.19 | 15.01±1.71 | 25.41±0.38 | 18.76±1.78 | 12.98±2.76 |
| PGL | 26.18±0.51 | 26.86±0.42 | 30.08±0.98 | 24.26±0.13 | 31.63±1.50 | 26.84±0.56 | 29.86±1.22 | 30.33±1.48 |
| PE | 40.01±1.07 | 41.59±1.21 | 44.87±2.06 | 43.56±1.81 | 49.49±2.68 | 42.93±1.95 | 48.59±2.62 | 52.72±3.60 |
| Lyso-PE | 3.36±1.46 | 4.21±1.50 | 2.33±1.66 | 3.61±1.96 | 3.78±2.33 | 4.75±2.06 | 2.78±1.91 | 3.87±2.17 |

SDS-PAGE analysis of LOS extracted from whole-cell lysates revealed no detectable differences in LOS abundance relative to wild-type (**Figure 3A**). All samples were normalized to OD_600_ 1.0 prior to extraction, and equal loading was verified using a specific antibody to the cytoplasmic protein RpoA. Densitometric quantification, showed that LOS levels remained statistically indistinguishable across all backgrounds, indicating that perturbations in PL trafficking or OM–PG tethering do not significant impact LOS biosynthesis or accumulation.

In contrast, the PL pool underwent clear and progressive remodeling (**Figure 3B**). Because these analyses were performed on total cellular lipid extracts and do resolve inner versus outer membrane lipid pools, the observed changes in PL composition are more consistent with perturbations in OM lipid organization. Using [³²P]-orthophosphate metabolic labeling followed by thin-layer chromatography (TLC), we quantified the relative abundance of the major PL species: phosphatidylethanolamine (PE), cardiolipin (CL), phosphatidylglycerol (PGL), and lyso-PE as previously described (12, 29). In wild-type cells, these species accounted for ∼40%, ∼26%, ∼30% and ∼3% of the total PL pool, respectively (**Table 1**).

Single mutants showed moderate but reproducible deviations from this profile (**Figure 3B**; **Table 1**). Δ*mla*, Δ*pldA* and Δ*tol-pal* mutants exhibited elevated PE levels (∼42–45%) and reduced CL (∼22–28%), while PGL remained largely stable except for a modest increase in Δ*pldA* (∼30%). These shifts were amplified in double mutants: Δ*mla* Δ*pldA*, Δ*mla* Δ*tol-pal* and Δ*pldA* Δ*tol-pal* strains accumulated 43-49% PE, whereas CL decreased to 15-25%. PGL increased slightly, but consistently, in all backgrounds lacking *pldA*. The triple mutant exhibited the most pronounced imbalance, with PE reaching 53%, CL dropping to 13%, and PGL rising to 30%.

Across the panel, the largest departures from the wild-type profile occurred in *pldA*-deficient backgrounds, which consistently showed stronger PE accumulation and deeper CL depletion than mutants lacking only *mla* or *tol-pal*. Notably, Δ*mla* displayed the greatest envelope fragility in stress assays (**Figure 2**), underscoring that Mla and PldA perturb distinct dimensions of OM homeostasis: Mla primarily preserves mechanical robustness, whereas PldA exerts dominant control over PL composition.

These data reveal a stepwise remodeling of the PL landscape as lipid homeostasis becomes progressively disrupted. The striking shifts in PE and CL, occurring despite stable LOS abundance, indicate that PL dysregulation, rather than LOS depletion, is the earliest and most defining biochemical signature associated with envelope destabilization in these mutants.

Together, these findings establish that dual disruption of PL transport and turnover creates a fragile but viable OM state, operationally defined as permissive. This state retains LOS yet exhibits altered PL abundance, increased permeability, and stress sensitivity, reducing the mechanical and compositional constraints that normally enforce glycolipid dependence.

### Disruption of lipid homeostasis reveals envelope stress-induced defects in growth, morphology, and cell division

To evaluate the physiological consequences of disrupting lipid homeostasis, we analyzed the growth and morphology of *A. baumannii* mutants under permissive and stress-inducing conditions. Standard LB broth provides a nutrient-rich, low osmolarity environment that buffers moderate defects in the envelope (30). In contrast, tryptic soy broth (TSB) imposes higher osmotic and metabolic demands, sensitizing cells with compromised OM architecture and revealing phenotypes that remain masked under permissive conditions (28).

In LB, all strains, including single (Δ*mla*, Δ*pldA*, Δ*tol*-*pal*), double mutants, and the triple mutant (Δ*ml*a Δ*pldA* Δ*tol*-*pal*) exhibited growth comparable to wild-type and reached similar optical densities (OD) at stationary phase (**Figure S3A**). These results indicate that under permissive conditions, the envelope remains functionally competent despite disruption of lipid transport and degradation pathways, suggesting that the cell can buffer moderate perturbations in OM lipid composition.

In contrast, growth in TSB revealed a graded hierarchy of envelope fragility (**Figure S3B**). Single mutants displayed mild delays during exponential growth, double mutants exhibited further attenuation and reduced final cell density, and the triple mutant showed the strongest phenotype, with a prolonged lag phase and markedly reduced yield. These progressive defects indicate that the combined disruption of lipid transport, lipid turnover, and OM tethering imposes an additive physiological burden that compromises the envelope’s ability to maintain integrity under osmotic/metabolic challenge.

To determine whether these growth defects reflected an underlying disorganization of the cell envelope, we analyzed cell morphology using phase-contrast microscopy and fluorescent D-amino acid (HADA) labeling to visualize sites of active PG synthesis. In LB, all strains maintained a coccobacillary-like morphology with clearly defined septa, consistent with proper envelope expansion and cell division (**Figure S3C**). Under TSB conditions, however, mutants exhibited striking abnormalities (**Figure S3D**): cells were elongated, frequently exhibited incomplete or mislocalized septa, and often formed short chains. These features are consistent with envelope stress and impaired coordination between the OM and the underlying PG layer.

Complementation assays confirmed the genetic basis of these phenotypes (**Figure S3E**). Expression of *mla*, *pldA*, or *tol*-*pal* in *trans* partially restored morphology in TSB, whereas empty vector controls had no effect. These results demonstrate that each system contributes to maintaining envelope integrity and proper cell division under osmotic/metabolic stress.

To test whether these phenotypes were driven by TSB-specific components or by osmotic imbalance itself, we exposed strains to LB lacking NaCl (LB_0_N), a defined hypoosmotic condition that increases turgor pressure while minimizing metabolic confounders (26). LB_0_N induced pronounced elongation and septation defects in all mutants, closely mirroring the morphological abnormalities observed in TSB (**Figure S3F**). Importantly, osmotic stabilization with sucrose restored wild-type morphology, eliminating the elongation and division defects across all backgrounds (**Figure S3G**). This osmo-protective effect confirms that the observed phenotypes arise primarily from mechanical stress imposed by hypoosmotic conditions rather than from metabolic limitations of the medium.

Collectively, these results highlight the critical role of lipid homeostasis in maintaining OM barrier function and coordinating OM expansion with PG synthesis. Disruption of PL transport, degradation, or OM tethering elicits a unified physiological response characterized by envelope instability, division defects, and hypersensitivity to environmental stress. These conditional vulnerabilities underscore the importance of lipid-envelope coordination in preserving cellular integrity under adverse conditions.

### Lipid asymmetry disruption enables emergence of colistin-resistant, LOS-deficient variants in *A. baumannii* strain ATCC 17978

We next asked whether this permissive architecture lowers the barrier for LOS loss under colistin selection and how cells remodel to stabilize a glycolipid-free envelope. Several *A. baumannii* strains, including AB5075 and ATCC 19606, can acquire colistin resistance through complete loss of LOS, typically via inactivation of early lipid A biosynthesis genes (12, 17, 31, 32). In contrast, the widely used reference strain ATCC 17978 consistently fails to generate LOS⁻ variants under standard colistin selection protocols (12), suggesting a rigid OM architecture with strict dependence on LOS for membrane integrity and viability.

Previous work has shown that deletion of *mrcA*, encoding the class A penicillin-binding protein PBP1A, partially relaxes this restriction and enables limited envelope remodeling (12, 13).

Building on this, we hypothesized that disrupting OM lipid homeostasis might similarly create a permissive state that lowers the threshold for LOS loss.

To test this, we measured colistin minimum inhibitory concentrations (MICs) using E-test strips (**Figure S4**). Wild-type strain ATCC 17978, as well as Δ*mla*, Δ*mla* Δ*pldA*, Δ*mla* Δ*tol*-*pal*, and the triple mutant Δ*mla* Δ*pldA* Δ*tol*-*pal*, all exhibited colistin MICs of 1 μg/mL, indicating no major change in baseline susceptibility. In contrast, Δ*pldA* showed a modest reduction (0.75 μg/mL), and Δ*tol*-*pal* and Δ*pldA* Δ*tol*-*pal* displayed slightly increased susceptibility (0.5 μg/mL). These small but reproducible shifts indicate that lipid-homeostasis defects can subtly modulate intrinsic susceptibility to colistin.

We next assessed whether these perturbations could facilitate the emergence of LOS-deficient variants by plating all strains on LB agar containing 10 μg/mL colistin, as established previously (12, 13, 17, 31, 32).

Strikingly, spontaneous colistin-resistant colonies consistently emerged only from the Δ*mla* Δ*pldA* background. SDS-PAGE analysis of whole-cell lysates confirmed LOS presence in wild-type and the parental Δ*mla* Δ*pldA* strain, whereas resistant isolates from this background completely lacked detectable LOS (**Figure 4A**). PL profiling of Δ*mla* Δ*pldA* LOS-deficient derivatives revealed no major additional remodeling relative to the permissive Δ*mla* Δ*pldA* background (**Figure S5A**), with only modest shifts in CL, PG, PE, and lyso-PE levels (**Figure S5B**), suggesting that the bulk of PL reorganization is already established in the permissive background prior to LOS loss. These results indicate that stable LOS deficiency is selectively accessible when both retrograde PL transport and surface PL degradation are simultaneously impaired under colistin selection. Complementation with *mla* and *pldA* abolished the emergence of LOS-deficient colonies, confirming that simultaneous disruption of PL transport and turnover enable LOS loss under colistin pressure (**Figure 4B**).

**Figure 4.**
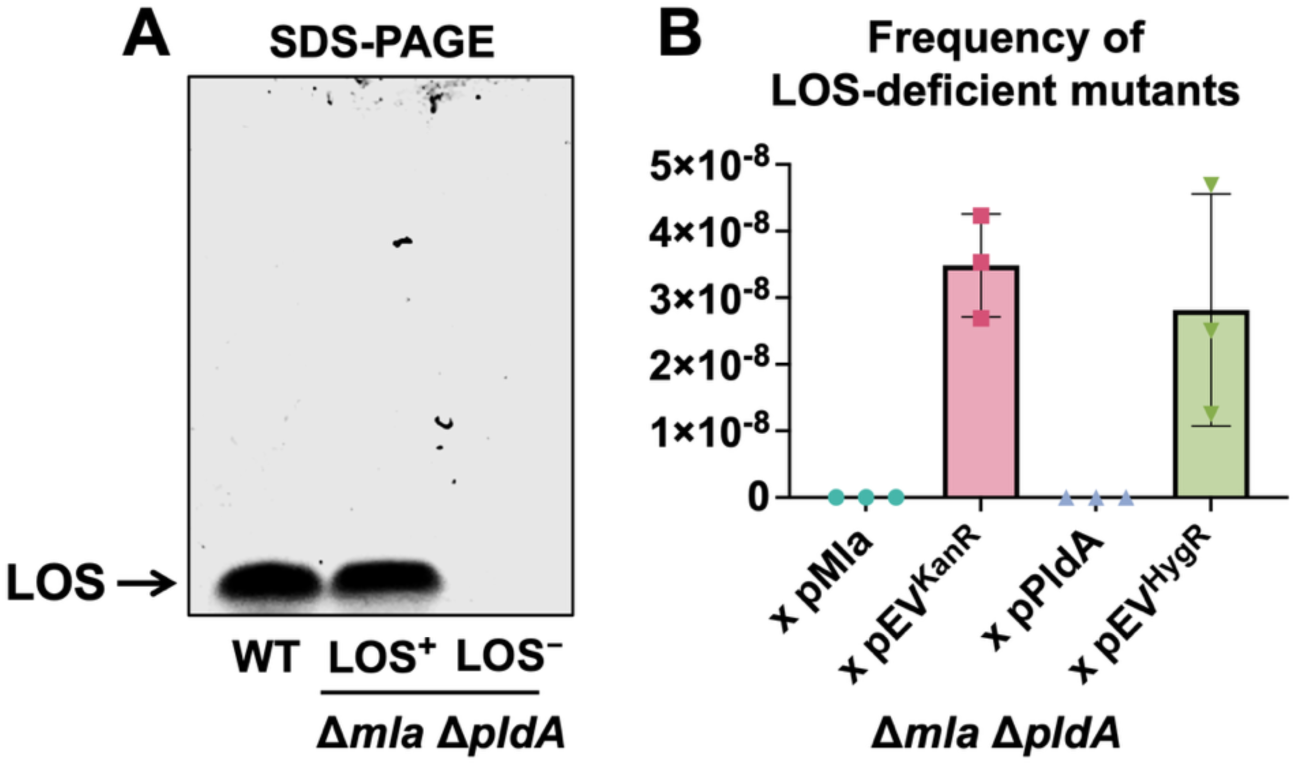
Disruption of PL transport and turnover enables emergence of LOS-deficient, colistin-resistant variants in *A. baumannii* strain ATCC 17978. (A) Colistin selection on LB agar (10 μg/mL) produced spontaneous resistant colonies exclusively from the Δ*mla* Δ*pldA* double mutant. SDS-PAGE analysis of whole-cell lysates followed by florescent staining shows that wild-type (WT) and the parental Δ*mla* Δ*pldA* strains retain LOS, whereas colistin-resistant isolates derived from this background lack detectable LOS entirely. (B) Genetic complementation of the Δ*mla* Δ*pldA* strain with plasmid-borne *mla* or *pldA* prevented the emergence of LOS-deficient variants under colistin selection, whereas empty vector controls retained the ability to generate LOS-deficient colonies. Error bars indicate standard deviation (*n* = 3).

Importantly, these LOS-deficient isolates correspond to the colistin-selected derivatives analyzed in subsequent experiments, enabling direct comparison across the same adaptive transition. This adaptation occurs independently of canonical lipid A modification pathways and instead reflects the attainment of an envelope-weakening threshold that permits selection and stabilization of LOS-deficient, colistin-resistant variants. Together, these results establish lipid asymmetry as a key barrier to LOS loss in strain ATCC 17978. Simultaneous disruption of retrograde PL transport and surface PL degradation thus unlocks an alternative, lipid-driven pathway to colistin resistance that remains inaccessible in the wild-type envelope architecture.

### Transcriptomic remodeling across three envelope states

To define how *A. baumannii* restructures its envelope during the transition from lipid asymmetry collapse to LOS loss and ultimately to colistin resistance, we compared the transcriptomes associated with three experimentally defined states derived from the Δ*mla* Δ*pldA* background following colistin selection. State 1 corresponds to Δ*mla* Δ*pldA* parental cells retaining LOS. State 2 represents LOS-deficient derivatives obtained after colistin selection and subsequently propagated in the absence of colistin. State 3 corresponds to LOS-deficient cells maintained under continuous colistin pressure. Each transition displayed a distinct transcriptional signature that together outline a coherent trajectory of envelope remodeling.

#### State 1: Lipid asymmetry collapse elicits a restrained and metabolically focused response

Inactivation of Mla and PldA triggered a limited transcriptional shift, with 67 differentially expressed genes relative to wild-type (**Figure 5A, Table S1**). Changes were largely confined to the affected loci, while most envelope systems, including LOS biosynthesis, lipoprotein trafficking, PG regulation, and multidrug efflux, remained near wild-type levels (**Figure 5D**). The dominant features were metabolic contraction, reflected by repression of phenylacetate catabolism, and surface simplification through downregulation of the *csu* adhesin operon. These patterns indicate that early defects in PL asymmetry are absorbed primarily through metabolic restraint rather than structural remodeling of the envelope, defining a primed but not yet adapted state.

**Figure 5.**
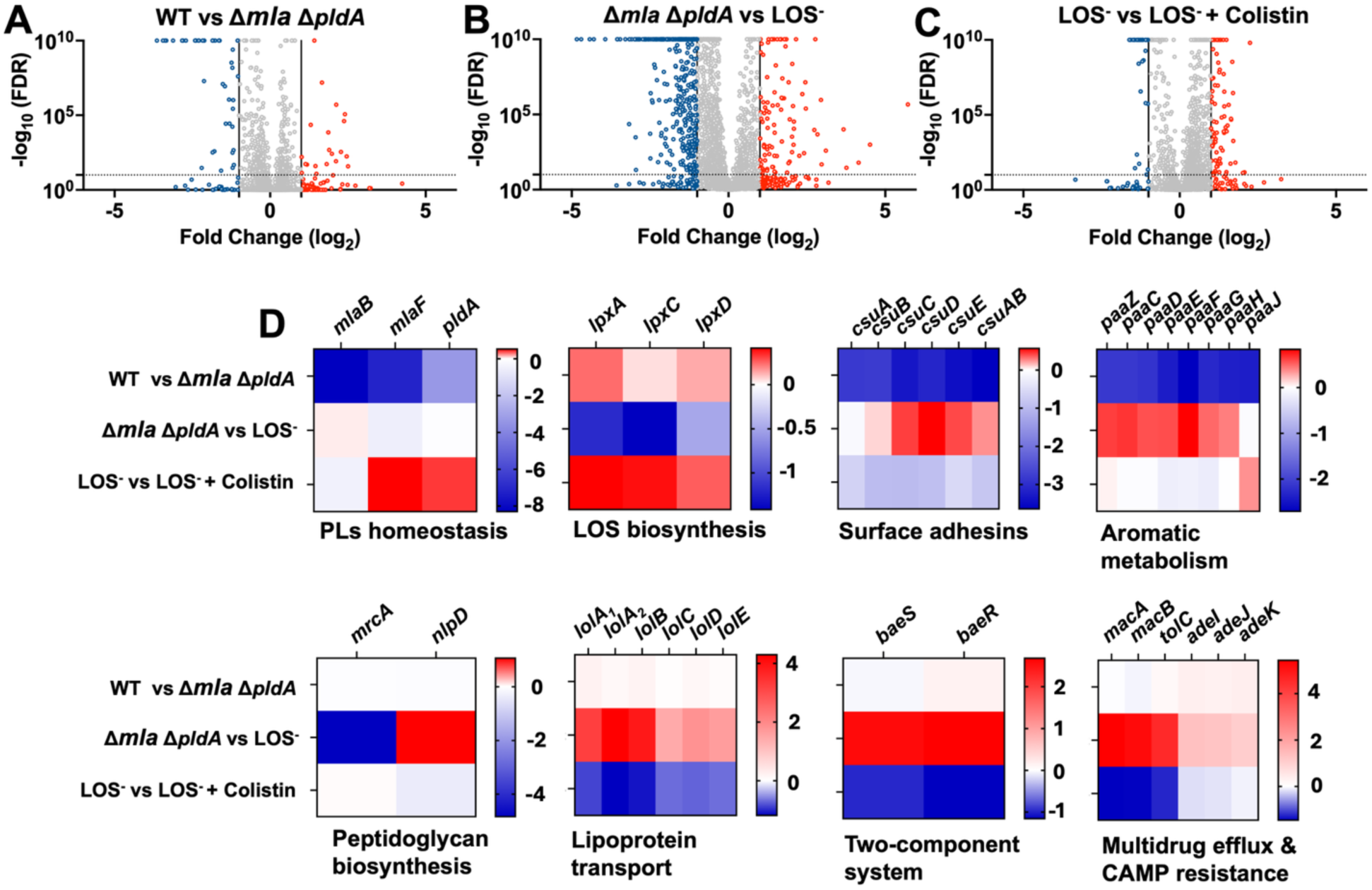
Transcriptomic reprogramming across envelope states from Δ*mla* Δ*pldA* to LOS-deficient, colistin-resistant states. (A-C) Volcano plots showing differential gene expression for the three sequential comparisons: (A) wild-type (WT) vs. Δ*mla* Δ*pldA* (State 1); (B) Δ*mla* Δ*pldA* vs. Δ*mla* Δ*pldA* LOS-deficient derivatives (State 1 vs State 2); and (C) Δ*mla* Δ*pldA* LOS-deficient cells propagated in the absence or presence of colistin (State 2 vs State 3). Data represent three independent biological replicates per condition. Significantly upregulated (red) and downregulated (blue) genes are defined by |log₂FC| ≥ 1, FDR < 0.05. (D) Heat maps summarizing expression patterns are key functional modules across the same comparation. These transcriptional trajectories outline a staged remodeling program, characterized by limited perturbations in Δ*mla* Δ*pldA*, extensive envelope and metabolic restructuring upon LOS loss, and focused consolidation under colistin pressure.

#### State 2: LOS deficiency triggers extensive envelope reorganization

The transition to a LOS-deficient state produced a major transcriptional reprogramming, with 629 differentially expressed genes (**Figure 5B, Table S1**). LOS loss induced strong activation of lipoprotein trafficking (LolA–E), the MacAB–TolC and AdeIJK efflux systems, and the BaeSR envelope stress regulators (**Figure 5D**). Early LOS biosynthesis genes (*lpxA*, *lpxC*, *lpxD*) were repressed, consistent with coordinated shutdown of lipid A production once LOS becomes dispensable. Surface adhesins, strongly repressed in State 1, were partially restored, indicating selective reactivation of surface functions during OM restructuring. PG regulation also shifted, with a pronounced decrease in *mrcA* (PBP1A) expression and modest induction of the septal regulator *nlpD* (**Figure 5D**).

Together, these changes define LOS loss as the principal trigger of a broad adaptive transcriptional program associated with envelope restructuring. Notably, this program builds upon a pre-existing envelope stress signature observed in the Δ*mla* Δ*pldA* background, with core responses such as lipoprotein trafficking, efflux systems, and stress signaling already initiated in State 1 and further expanded upon transition to the LOS-deficient state. This expansion includes additional modulation of outer membrane proteins, lipoproteins, and PG-associated factors, consistent with active remodeling of envelope architecture.

#### State 3: Colistin exposure fine-tunes the LOS-deficient envelope

Colistin treatment elicited a more focused and comparatively mild transcriptional response (111 differentially expressed genes; **Figure 5C, Table S1**). Pathways strongly activated during LOS loss, including lipoprotein trafficking and BaeSR signaling, declined toward intermediate levels (**Figure 5D**). Efflux systems did not intensify; instead, MacAB–TolC and AdeIJK receded toward baseline, indicating that their induction is driven by LOS deficiency rather than by colistin itself. The *csu* operon, partially restored in State 2, became downregulated again under colistin, consistent with further simplification of the cell surface. These observations show that colistin does not globally amplify the LOS-loss program but refines it as the LOS-free envelope stabilizes under antibiotic pressure.

Together, these transitions reveal an ordered remodeling trajectory in which *A. baumannii* first buffers lipid asymmetry collapse through metabolic restraint (a primed state), then undergoes a decisive restructuring of envelope architecture upon LOS loss (an adaptive state), and finally fine-tunes this rebuilt envelope under colistin pressure. This progression indicates that colistin-resistant growth without LOS emerges not from a single shift, but from a sequential traversal of discrete envelope states that progressively recalibrate membrane structure and function.

### Reduced PBP1A levels are required for the transition to LOS deficiency

Transcriptomic analysis identified *mrcA*, which encodes the bifunctional PG synthase PBP1A, as one of the most strongly repressed genes in LOS-deficient cells (**Figure 5D**). Because previous studies showed that *mrcA* inactivation alone can generate LOS-deficient variants in *A. baumannii* ATCC 17978 (12, 13), we asked whether reduced PBP1A abundance is a defining feature of the LOS⁻ state.

To determine whether this transcriptional pattern corresponds to changes at the protein level, we next assessed PBP1A abundance in the same Δ*mla* Δ*pldA*-derived LOS-deficient isolates described above (**Figure 6A**). Wild-type and Δ*mla* Δ*pldA* cells exhibited comparable PBP1A levels, indicating that collapse lipid asymmetry alone does not alter the steady-state abundance of this enzyme. In contrast, LOS-deficient derivatives of the Δ*mla* Δ*pldA* background displayed a marked reduction in PBP1A, demonstrating that loss of this protein coincides specifically with entry into the LOS⁻ state. As expected, Δ*mrcA* and Δ*mrcA* LOS⁻ strains lacked detectable PBP1A. We next tested whether preventing this decline in PBP1A was sufficient to block the transition to LOS deficiency (**Figure 6B**). In the Δ*mla* Δ*pldA* background, empty-vector controls produced LOS-deficient colonies at frequencies of ∼10⁻⁸. In contrast, expression of *mrcA* from either its native promoter or an IPTG-inducible promoter abolished detectable LOS⁻ variants, indicating that maintaining PBP1A availablity restricts access to the LOS-deficient state rather than merely reducing its frequency.

**Figure 6.**
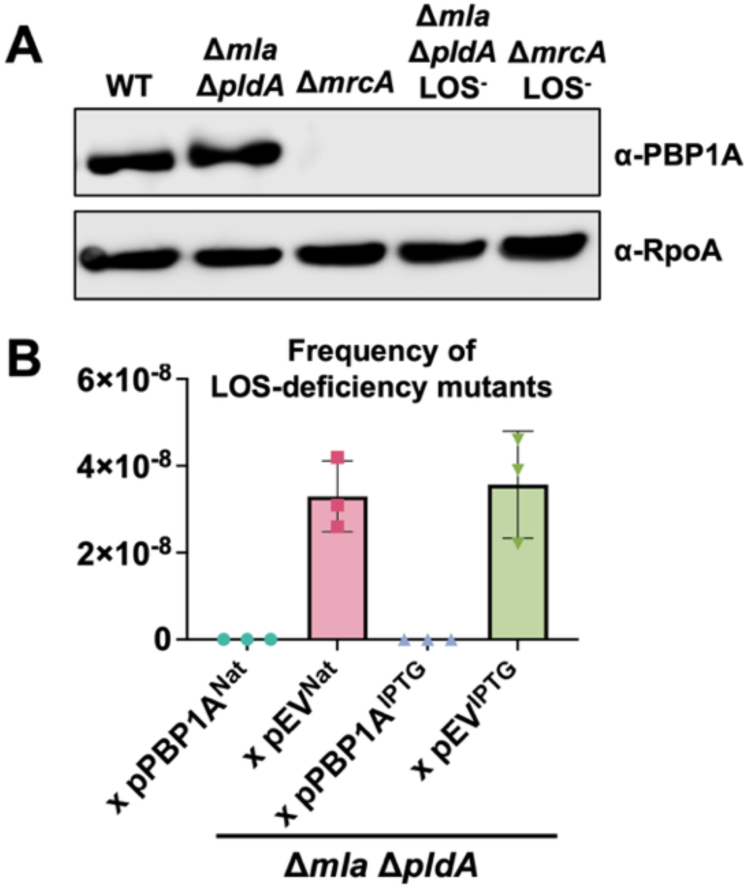
Reduced PBP1A facilitates the transition to LOS deficiency. (A) Immunoblot analysis of PBP1A abundance in wild-type (WT), Δ*mla* Δ*pldA*, Δ*mrcA*, LOS-deficient derivatives of the Δ*mla* Δ*pldA* background described in Figure 4. PBP1A (94.74 kDa) is detected in WT and the Δ*mla* Δ*pldA* double mutant, but is absent in Δ*mrcA* and markedly reduced in LOS-deficient isolates. RpoA (37.62 kDa) is shown as a loading control. (B) Sustained PBP1A expression prevent LOS loss. Expression of *mrcA* (PBP1A) in the Δ*mla* Δ*pldA* background, either from a native-promoter construct (Nat) or an IPTG-inducible plasmid (IPTG), blocks the emergence of LOS-deficient, colistin-resistant colonies. In contrast, empty-vector controls, in contrast, readily yield LOS-deficient variants. These results demonstrate that PBP1A availability is a critical determinant governing the transition of Δ*mla* Δ*pldA* cells can transition to a stable LOS-deficient state. Error bars indicate standard deviation (*n* = 3).

Together, these findings show that reduced PBP1A abundance is not simply correlated with LOS loss but is functionally required for it. Δ*mla* Δ*pldA* cells retain PBP1A at wild-type levels, indicating that lipid asymmetry disruption alone is insufficient to diminish this enzyme. Instead, PBP1A levels decline specifically upon transition to the LOS⁻ state, and genetic or plasmid-based maintenance of *mrcA* prevents this transition. Conversely, complete inactivation of PBP1A through *mrcA* deletion provides an alternative route to the same outcome. These results establish that LOS and PBP1A reduction are tightly coupled features of the same adaptive transition, rather than independent events. Together, these convergent paths point to a common architectural principle: reducing PBP1A-mediated PG expansion is essential for stabilizing a LOS-free envelope.

### Synergistic disruption of PG synthesis and lipid asymmetry expands access to LOS deficiency in *A. baumannii*

Having established that reduced PBP1A abundance is required for the LOS⁻ state, we next asked whether weakening PG synthesis actively cooperates with disrupted lipid asymmetry to reshape envelope stability. This question follows naturally from our earlier observation: although *mrcA* expression decreases only after cells become LOS⁻, genetic inactivation of *mrcA* alone is sufficient to permit LOS deficiency (12, 13). We therefore constructed a triple mutant lacking *mrcA* in the Δ*mla* Δ*pldA* background to directly test whether removing PBP1A expands the structural window in which LOS loss becomes accessible.

Proliferation in rich medium was comparable across wild-type, Δ*mla* Δ*pldA*, Δ*mrcA*, and the Δ*mla* Δ*pldA* Δ*mrcA* triple mutant, indicating that these mutations do not impair growth and viability under non-stress conditions (**Figure 7A**). However, phase-contrast imaging and HADA labeling revealed distinct morphological abnormalities (**Figure 7B**). Whereas Δ*mla* Δ*pldA* cells retained near–wild-type morphology, both Δ*mrcA* and the triple mutant displayed pronounced elongation and septation defects, consistent with disrupted coordination between elongation and division.

**Figure 7.**
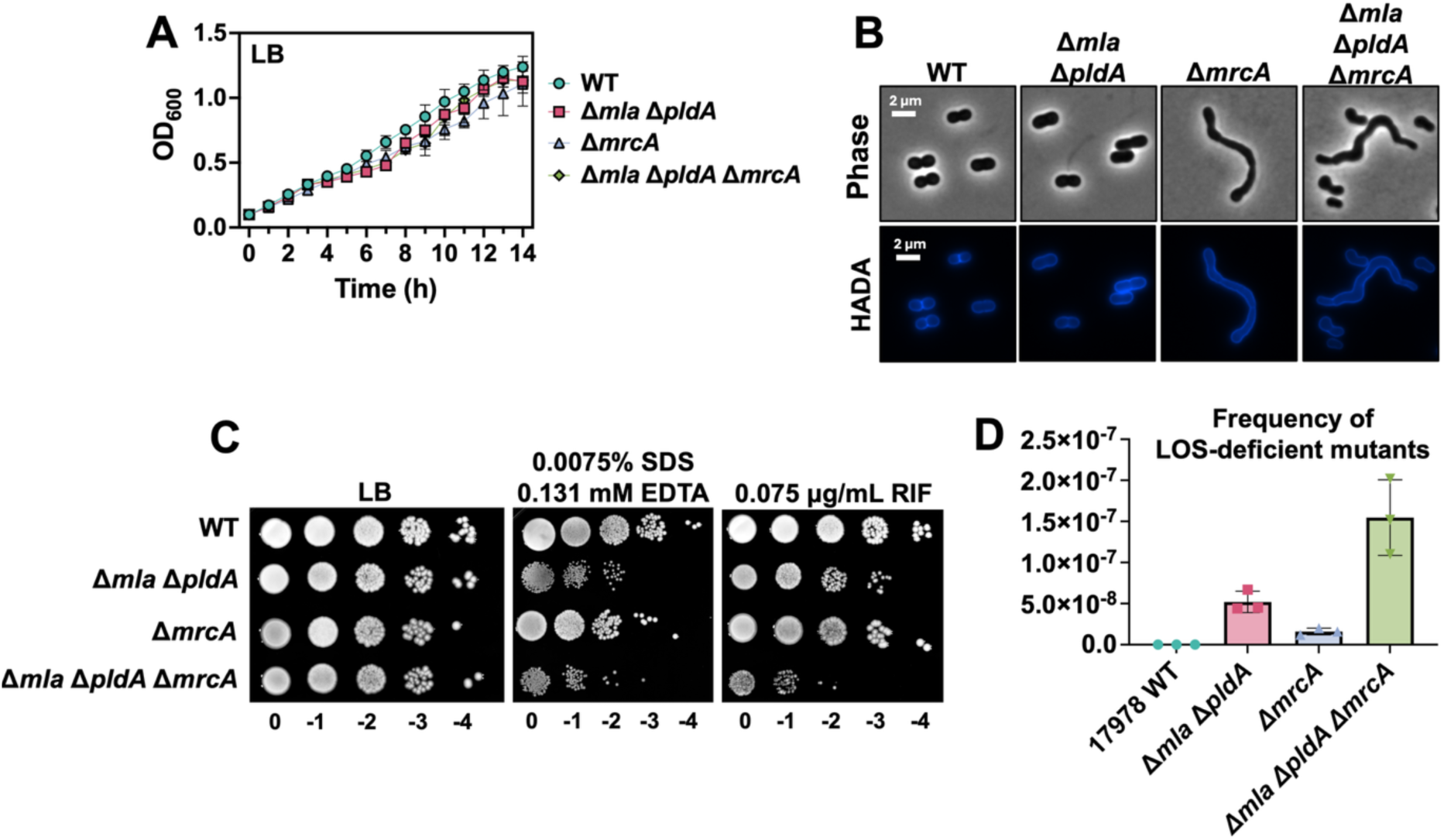
Disruption of peptidoglycan synthesis exacerbates envelope stress and promotes LOS loss in the Δ*mla* Δ*pldA* background. (A) Growth curves in LB medium show comparable proliferation among wild-type (WT), Δ*mla* Δ*pldA*, Δ*mrcA*, and the triple mutant Δ*mla* Δ*pldA* Δ*mrcA*, indicating that combined envelope perturbations do not impair basal growth under non-stress conditions. (B) Phase-contrast and HADA fluorescence microscopy reveal morphological alterations in Δ*mrcA* and the triple mutant, including cell elongation and defects in septation, hallmarks of impaired PG synthesis due to loss of PBP1A activity. In contrast, Δ*mla* Δ*pldA* cells retain WT-like morphology. Scale bars: 2 µm. (C) Quantification of colony-forming units following treatment with mild detergent (0.0075% SDS, 0.131 mM EDTA) or subinhibitory rifampicin (0.075 µg/mL) reveals synthetic higher sensitivity in the triple mutant, reflecting compounded defects in OM permeability and barrier integrity. (D) Frequency of LOS-deficient variant emergence under colistin selection (10 μg/mL). While Δ*mla* Δ*pldA* and Δ*mrcA* strains yield resistant colonies, the triple mutant exhibits a more than twofold increase, indicating that combined disruption of lipid and PG homeostasis synergistically promotes LOS loss and resistance acquisition. Error bars indicate standard deviation (*n* = 3).

OM integrity assays further highlighted the interaction between these pathways (**Figure 7C**). Δ*mla* Δ*pldA* and Δ*mrcA* mutants showed moderate susceptibility to mild detergent stress and to subinhibitory rifampicin, whereas the triple mutant was substantially more sensitive, reflecting additive compromise of envelope barrier function.

We then quantified the frequency of LOS-deficient variants under colistin selection (**Figure 7D**). No resistant colonies emerged from wild-type cultures. LOS-deficient derivatives appeared from Δ*mla* Δ*pldA* and Δ*mrcA* strains at frequencies of approximately 4–7×10⁻⁸ and 1.2–2×10⁻⁸, respectively. Strikingly, the triple mutant produced LOS-deficient, colistin-resistant colonies at markedly higher frequencies (1.1–2.0×10⁻⁷). These results indicate that simultaneous disruption of PG synthesis and PL homeostasis produces a permissive envelope state that greatly enhances access to the LOS⁻ architecture.

Together, these findings demonstrate that PG synthesis and lipid asymmetry maintenance are interdependent axes of OM stability. Their combined disruption amplifies envelope destabilization and expands the adaptive landscape, enabling evolutionary trajectories that remain inaccessible in the wild-type. This synergy highlights the architectural plasticity of *A. baumannii* and reveals vulnerabilities that may be exploited to restrict the emergence of LOS-deficient, colistin-resistant states.

### LOS-deficient variants exhibit a trade-off between colistin resistance and envelope robustness

These hierarchical adaptations suggested that each transcriptional state carries distinct physiological consequences. To test these predictions, we characterized LOS-deficient variants arising under colistin selection from the Δ*mla* Δ*pldA* background and compared them to both their parental mutant and the wild-type *A. baumannii* ATCC 17978 under various stress conditions.

In rich LB, LOS⁻ variants exhibited markedly reduced growth compared to both wild-type and the Δ*mla* Δ*pldA* strain, indicating a significant fitness cost associated with LOS absence under non-selective conditions (**Figure S6A**). However, in the presence of 10 μg/mL colistin, a concentration lethal to both wild-type and parental mutant, the LOS⁻ variant grew robustly. This demonstrates a clear physiological trade-off: while LOS loss compromises baseline fitness, it provides a pronounced survival advantage under colistin pressure.

MICs analysis by E-test revealed a striking inversion in antibiotic susceptibility profiles (**Figure S6B**). The Δ*mla* Δ*pldA* strain remained susceptible to colistin and exhibited elevated MICs to vancomycin, daptomycin, and bacitracin. By contrast, the LOS⁻ variants displayed increased MICs to colistin but decreased MICs to these antibiotics, consistent with the dual consequences of LOS loss on lipid A-mediated targeting and OM barrier function.

Morphological analysis revealed severe envelope defects in LOS⁻ variants (32) (**Figure S6C**). Whereas wild-type and Δ*mla* Δ*pldA* cells maintained a coccobacillary morphology with defined septa, LOS-deficient strains formed elongated or curved cells, exhibited irregular septation, and frequently aggregated. These defects were consistently observed in LOS-deficient derivatives arising from both the Δ*mrcA* single mutant and the Δ*mla* Δ*pldA* Δ*mrcA* triple mutant background (**Figure S6D**), indicating that transition to the LOS-deficient state is associated with pronounced morphological alterations, independent of the genetic background.

Intriguingly, exposure to colistin partially restored morphology in LOS-deficient cells, reducing aggregation and improving septation. This paradoxical effect may reflect transient stabilization of the disorganized OM upon colistin binding to surface-exposed PLs.

Together, these findings define a clear physiological trade-off: acquisition of colistin resistance through LOS loss is accompanied by compromised OM robustness and heightened vulnerability to other antibiotics. This balance illustrates how envelope remodeling enables resistance by reshaping membrane architecture, while simultaneously exposing vulnerabilities that constrain evolutionary trajectories and reveal collateral sensitivities

## Discussion

The Gram-negative OM achieves barrier integrity through strict lipid asymmetry, with PLs confined to the inner leaflet and LPS/LOS forming the outer leaflet. Our data support a model in which lipid asymmetry functions as a structural checkpoint that governs access to glycolipid-independent survival and thereby shapes evolutionary routes to colistin resistance (**Figure 1**). When quality-control systems are intact, strain ATCC 17978 remains strictly LOS-dependent and does not traverse LOS loss under selection. In contrast, deletion of retrograde PL transport (Mla) and surface PL turnover (PldA) collapses asymmetry, creating a fragile but permissive envelope (**Figure 2**) with increased detergent and rifampicin sensitivity, thereby lowering the barrier to LOS loss.

Integrated genetics, lipidomics, and transcriptomics define a general three-state trajectory—basal to permissive to adapted (**Figure 1**): (1) In the basal state (LOS-dependent), lipid asymmetry is actively maintained, enforcing glycolipid essentiality and ensuring low OM permeability, with colistin targeting LPS/LOS. (2) The permissive state (asymmetry collapsed) emerges when PL homeostasis is disrupted through loss of Mla and PldA, producing a metastable OM that retains LOS yet exhibits heightened stress sensitivity (**Figure 2**). Lipidomics analyses revealed PE accumulation and CL depletion across mutant combinations (**Figure 3**; **Table 1**), most pronounced in *pldA*-deficient backgrounds. While CL depletion is consistently associated with these perturbations, it is not sufficient to define the permissive state. Because these analyses were performed on total cellular extracts, membrane fractionation approaches (33) provide a framework to resolve lipid distributions between inner and outer membranes. Within this context, the observed PL remodeling is most consistent with perturbations in OM lipid organization, although membrane-specific distributions cannot be directly assigned from our data. (3) Finally, in the adapted (LOS-deficient) state, cells rebuild the OM around protein/lipoprotein scaffolds, attenuate PG expansion through reduced PBP1A levels, and stress response pathways (**Figure 5**), enabling colistin resistant growth while introducing collateral vulnerabilities.

These results reveal two interdependent axes that together determine envelope viability. The first axis is lipid asymmetry, maintained by Mla and PldA. When PL transport and turnover are destabilized, outer-leaflet PLs (notably PE) accumulate, weakening mechanical cohesion and exposing the OM to SDS, EDTA, and rifampicin stress (**Figure 2**). Despite unchanged LOS abundance (**Figure 3A**), the PL pool undergoes stepwise remodeling (**Figure 3B**; **Table 1**). Importantly, not all mutant backgrounds displaying similar shifts in PL composition exhibit the same capacity to generate LOS-deficient variants, underscoring that bulk lipid composition alone does not define the permissive state. Instead, disruption of PL trafficking and membrane organization emerges as a key determinant of whether cells can access this state. The second axis is PG synthesis, mediated by PBP1A. Transition to LOS deficiency consistently coincides with sharp PBP1A repression, and maintaining PBP1A activity blocks adaptation to LOS loss (**Figure 6**). Functionally, decreasing PG synthetic pressure widens the mechanical tolerance window of a destabilized OM, as demonstrated by the Δ*mla* Δ*pldA* Δ*mrcA* triple mutant, which exhibits the highest stress susceptibility and the greatest frequency of LOS⁻ variant emergence frequency (**Figure 7**). This synergy explains how cells traverse the permissive gate and stabilize a glycolipid-free envelope: collapse asymmetry to reduce constraints, then throttle PG expansion to minimize envelope mismatch.

Our engineered progression mirrors naturally evolved suppressors seen in LOS-deficient *A. baumannii* lineages, where mutations in *mla* and *pldA* frequently arise (22, 32). These lesions weaken PL quality control, enabling outer-leaflet PL retention that compensates for absent glycolipids and stabilizes alternate OM configurations (**Figures 3 and 5**). Likewise, PBP1A dosage functions as an evolutionary gate: low basal PBP1A pre-primes LOS-free survival, whereas elevating PBP1A closes the gate (**Figure 6**). In ATCC 17978, entry into the LOS-deficient state requires reduced PBP1A abundance, and sustained PBP1A blocks LOS loss (12). Together, these axes explain heterogeneous LOS essentiality across lineages and predict convergent routes by which strains relax glycolipid dependence under antibiotic pressure.

Although dissected in *A. baumannii*, the underlying principles are species-agnostic: retrograde PL trafficking, outer-leaflet PL clearance, and OM–PG coupling are conserved features of Gram-negative envelope homeostasis. Reports of conditional LPS/LOS dispensability in diverse taxa (9–13, 17, 31) align with our architecture-first model (**Figure 1**), which posits lipid asymmetry as a gate and PG synthesis as a throttle—design rules likely relevant under polymyxin pressure or LPS/LOS biosynthesis inhibition.

If asymmetry collapse is the gate to LOS-independent survival, then preserving Mla/PldA function—or mimicking their activities—may help maintain glycolipid dependence and restrict access to LOS-deficient trajectories (**Figures 2–4**). Conversely, metabolic states that enrich PE or deplete CL may promote the permissive state (**Figure 3**; **Table 1**). On the PG axis, sustaining PBP1A activity (or countering its repression) appears to close the LOS-loss gate (**Figure 6**); pairing polymyxins or LPS/LOS-biosynthesis inhibitors with agents that increase PG synthetic pressure may harden the envelope against glycolipid-free remodeling (**Figure 7**). Finally, the collateral sensitivities of LOS-deficient cells (susceptibility to large hydrophilic antibiotics; morphology defects) offer combination opportunities (**Figure S5**).

The Tol-Pal contribution to PL organization and OM mechanics in *A. baumannii* merits deeper dissection (**Figure 2; Figure S1**). Testing the model across clinical isolates, varied metabolic states, and *in vivo* contexts will quantify how often permissive–adapted transitions occur, and which suppressor paths dominate. Defining biophysical thresholds (e.g., PE:CL ratios, PG synthesis rates) should enable predictive models of resistance trajectories under different regimens.

By uncovering lipid asymmetry as a structural checkpoint and PBP1A-mediated PG synthesis as a mechanical throttle, we provide a framework to explain how Gram-negative envelopes transition from LOS-dependent to glycolipid-free survival under colistin pressure (**Figure 1**). The three-state model integrates recurrent suppressors in *mla**/**pldA* and *mrcA* dosage effects to account for strain-specific differences in LOS essentiality and to predict convergence on LOS-independent resistance.

## STAR★Methods

### Key resources table

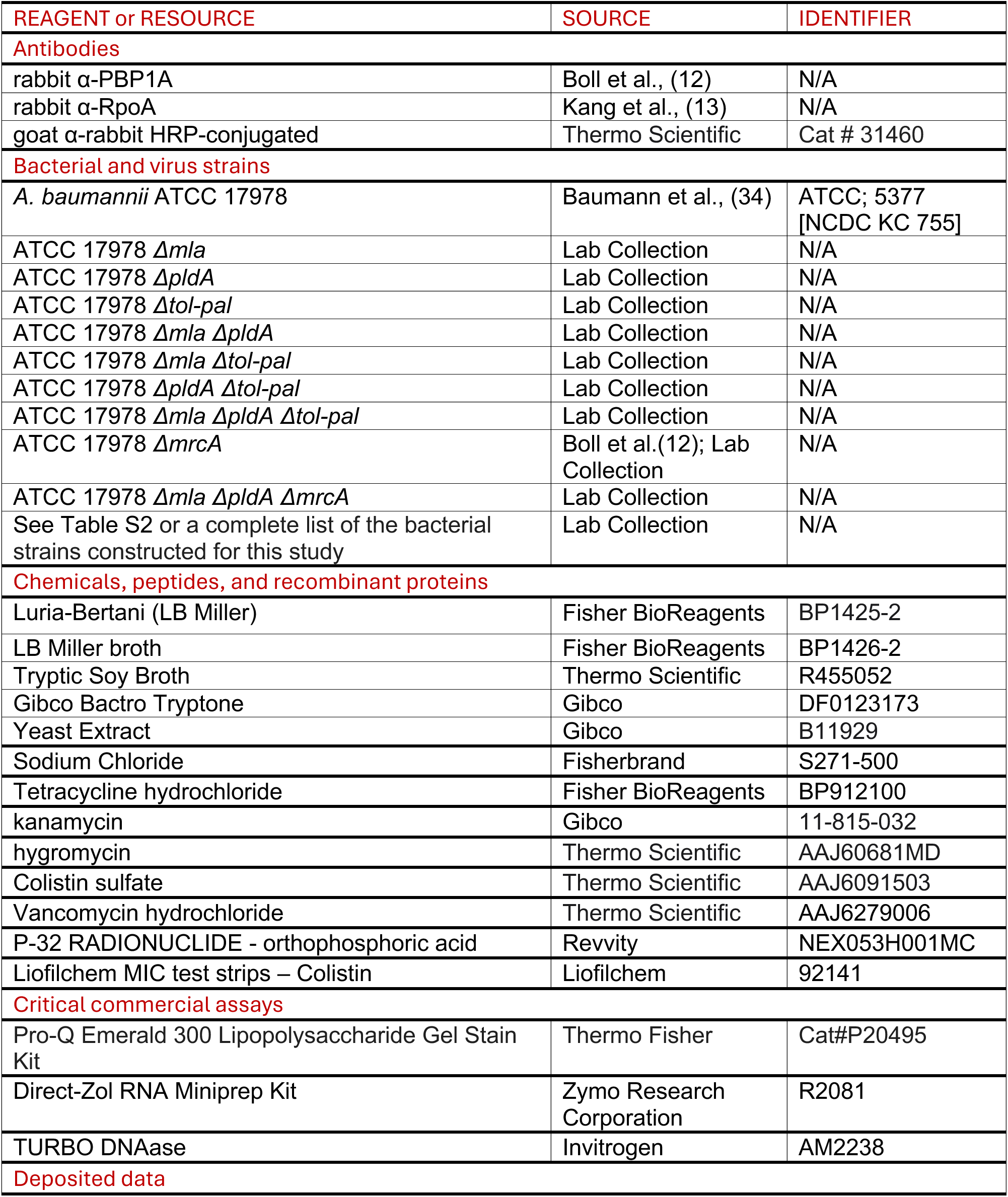
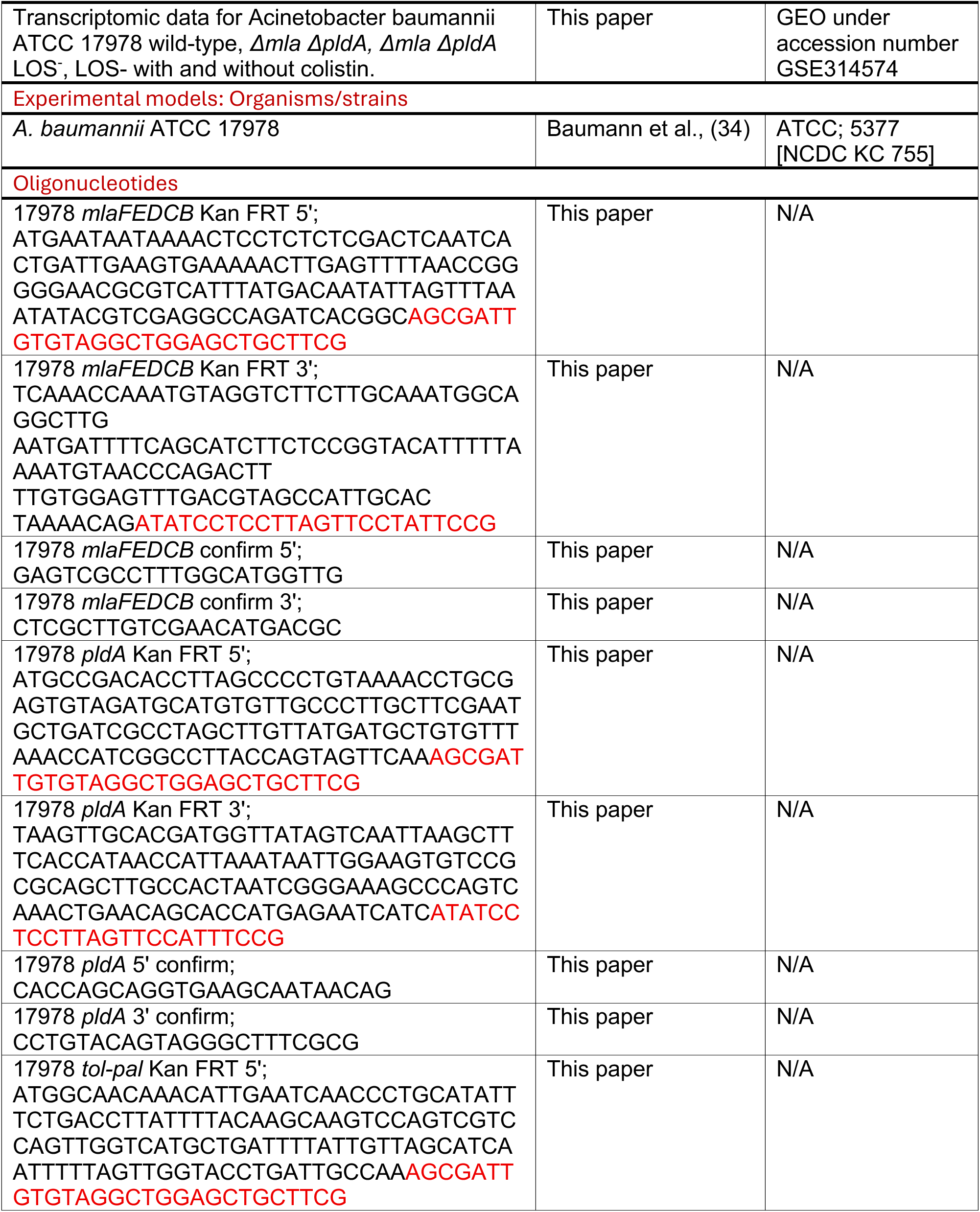
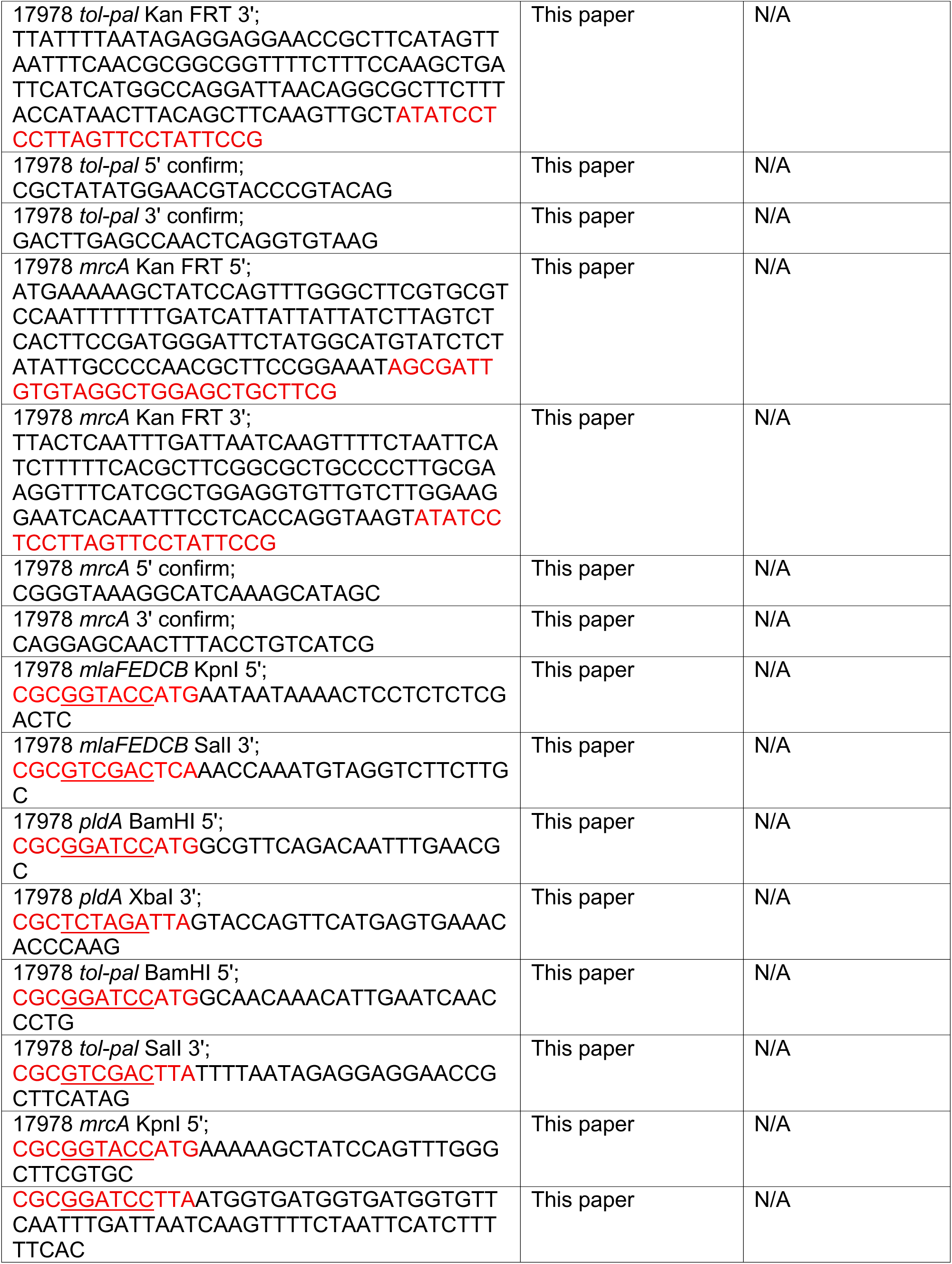
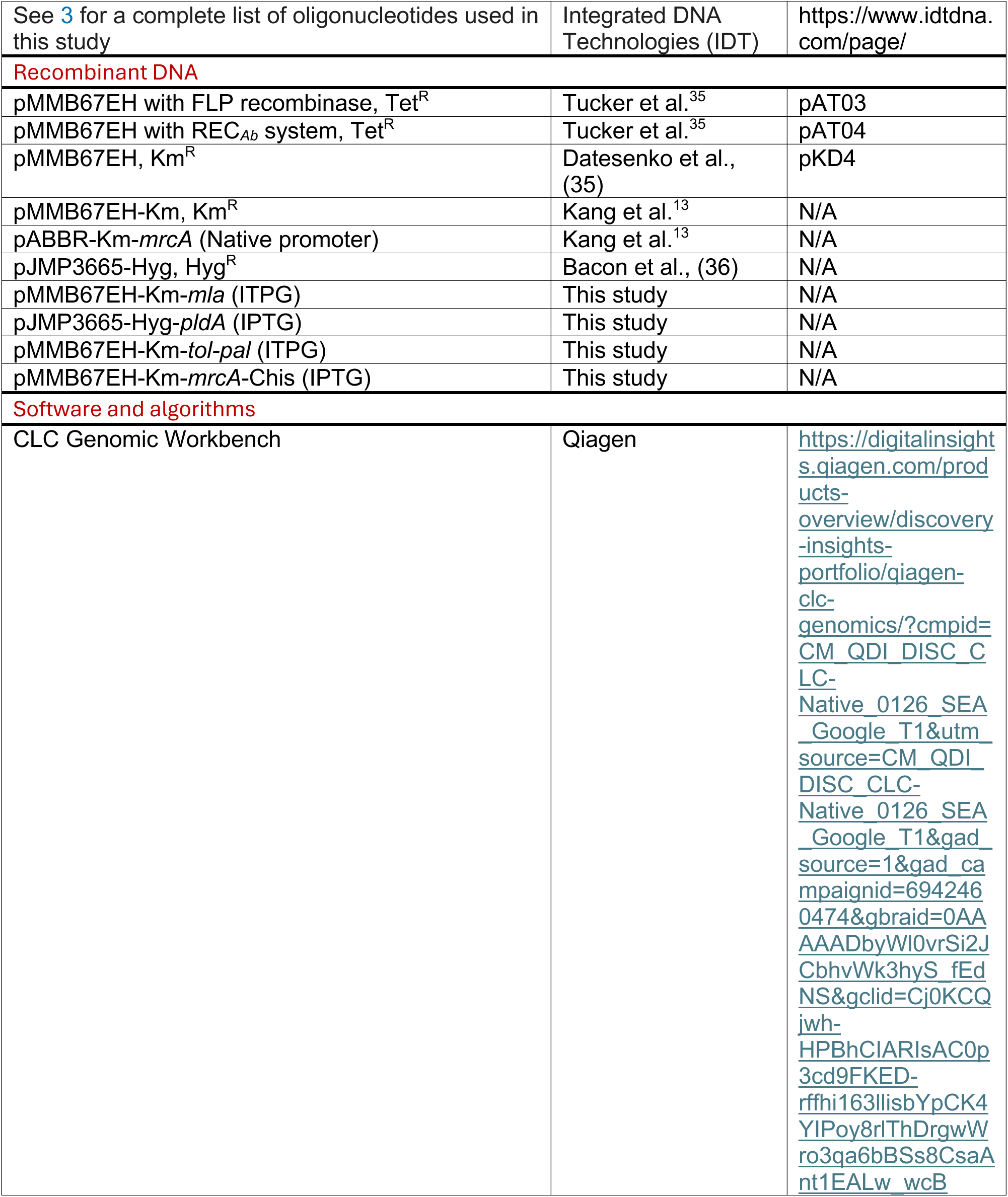
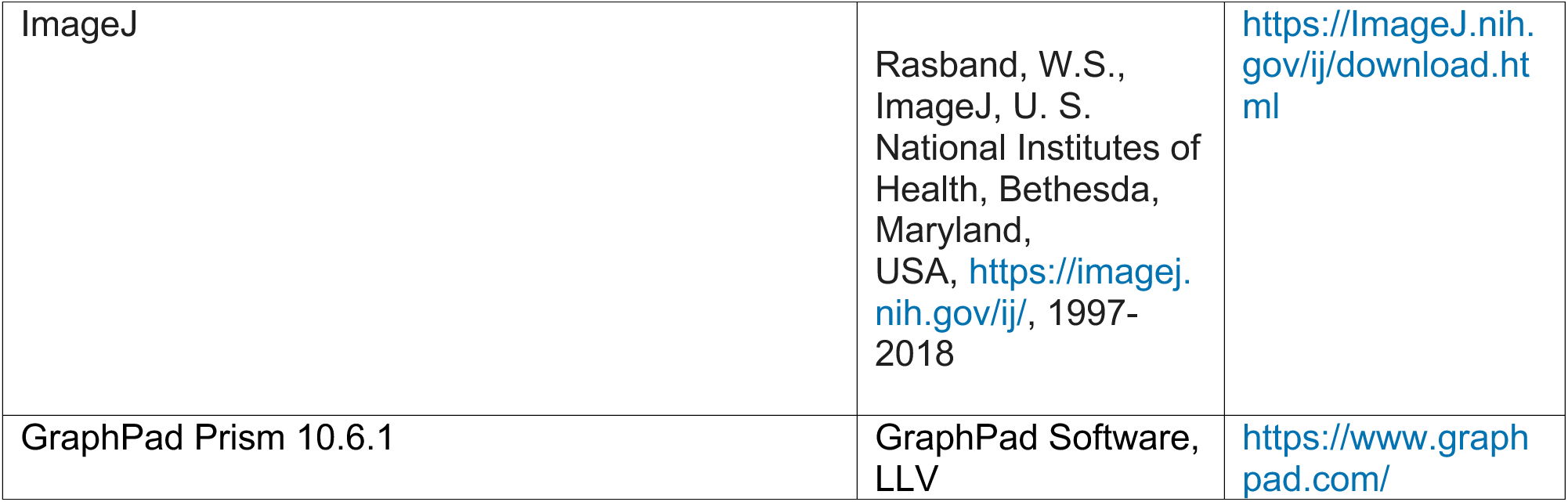

### Resource Availability Lead Contact

Further information and requests for resources and reagents should be directed to and will be fulfilled by the lead contact, Joseph M. Boll.

## Materials Availability

All unique/stable reagents generated in this study are available from the lead contact upon request, but we may require a payment and/or a materials transfer agreement if there is potential for commercial application.

### Experimental Model and Subject Details

All *Acinetoabcter baumannii* strains were derived from strain ATCC 17978 (34) and are listed in Table S2, and primer sequences are provided in Table S3. *A. baumannii* ATCC 17978 was routinely cultured aerobically at 37°C on Luria-Bertani (LB Miller) agar or in LB Miller broth, Tryptic Soy Broth (TSB) or LB without added NaCl (LB_0_N). Liquid cultures were grown with shaking unless otherwise indicated.

Where indicated, antibiotics were used at the following final concentrations: tetracycline (10 μg/mL), kanamycin (15 or 25 μg/mL), hygromycin (250 μg/mL), colistin (10 μg/mL), and vancomycin (10 μg/mL).

ATCC 17978 is known to comprise two variants (37). Colony PCR using primers specific to the CL synthase gene (*clsC2*) confirmed the presence of a PCR product, indicating that experiments were performed in the 17978 UN variant.

## Methods Details

### Construction of mutant and complemented *A. baumannii* strains

*A. baumannii* mutants were generated using established Rec*_Ab_*-based recombineering protocols (29, 38–40). Briefly, strains carrying the pMMB67EH^TetR^ plasmid encoding REC*_Ab_* were diluted from overnight cultures into 50 mL LB containing tetracycline (10 µg/mL) to an initial OD_600_ 0.05 and incubated for 40 minutes. Expression of REC*_Ab_* was induced with IPTG (2 mM), and cultures were grown at 37 °C until mid-log phase (OD_600_ 0.4). Cells were washed five times with ice-cold 10% glycerol and concentrated. Approximately 10¹⁰ cells were electroporated in a 2-mm cuvette at 1.8 kV with 5 µg of linear PCR product. After electroporation, cells were recovered in 5 mL LB containing IPTG (2 mM) for 4 h and plated on LB agar containing kanamycin (15 µg/mL). Candidate mutants were verified by PCR.

To cure the pMMB67EH^TetR^::REC*_Ab_* plasmid after mutant isolation, strains were streaked for single colonies on LB agar containing NiCl₂ (2 mM), as described (29, 38–41). Cured insertion mutants were transformed with pMMB67EH^TetR^ encoding FLP recombinase, recovered for 1 h in LB, and plated on LB agar containing tetracycline (10 µg/mL) and IPTG (2mM) to induce FLP. Excision of the kanamycin cassette was verified by PCR. Double and triple mutants were constructed sequentially using the same workflow.

For genetic complementation, the *mlaF*-*B* coding region (*A1S_3103*–*A1S_3099*) was cloned into the KpnI/SalI sites, and *tolQ*-*pal* (*A1S_2591*–*A1S_2595*) was cloned into the BamHI/SalI sites of pMMB67EH^KanR^. The *pldA* coding sequence (*A1S*_*1919*) was cloned into the BamHI/XbaI sites of pJMP3665^HygR^.

To probe the interaction between PG synthesis and OM lipid homeostasis, *mrcA* (*A1S_3197*) was cloned into pMMB67EH^KanR^ using KpnI/BamHI restriction sites and expressed *in trans* in the Δ*mla* Δ*pldA* background.

Plasmids were introduced into the indicated strains, and cultures were maintained under appropriate antibiotic selection (25 µg/mL kanamycin for pMMB67EH^KanR^ and 250 µg/mL hygromycin for pJMP3665^HygR^). Gene expression was induced with IPTG (1 mM).

### SDS/EDTA and antibiotic susceptibility assays

Overnight cultures were diluted to OD_600_ 0.05 in fresh LB. Serial dilutions (10⁰ to 10⁻⁴) were prepared and spotted onto LB agar plates containing the indicated concentrations of SDS/EDTA or rifampicin. Plates were incubated at 37 °C, and growth was scored after incubation.

### Analysis of ³²P-labeled PLs

PL composition was analyzed as described (12, 29, 42) with minor modifications. Overnight cultures were diluted to OD_600_ 0.05 in 5 mL LB containing 5 µCi/mL [³²P]-orthophosphate (PerkinElmer) and grown at 37 °C with shaking for 3 h. Cells were then harvested, and total lipids were extracted using the Bligh and Dyer method (43).

Radiolabeled PLs were separated by thin-layer chromatography (TLC) on silica gel 60 plates using chloroform, methanol, and acetic acid (60:25:5, v/v). Plates were dried, exposed to a phosphorimaging screen, and scanned using an Amersham Typhoon FLA 9500 (GE Healthcare). Bands intensities were quantified in ImageJ, and each lipid species was expressed as a percentage of the total radiolabeled signal per lane.

### LOS staining and analysis

LOS profiles were assessed as described (29, 44) with modifications. Overnight cultures were diluted to OD_600_ 0.05 in 5 mL LB and grown at 37 °C with shaking to OD_600_ 1.0. Cells were pelleted (13,000 rpm, 5 minutes) and resuspended in 1× LDS sample Buffer supplemented with β-mercaptoethanol, then boiled for 10 minutes. After cooling, proteinase K was added, and samples were incubated at 55 °C overnight. Samples were boiled for 5 minutes and resolved by SDS-PAGE. Gels were fixed and stained using Pro-Q Emerald 300 LPS Gel Stain Kit (Thermo Fisher Scientific, P20495) according to the manufacturer’s instructions.

### Growth curves and cell morphology

Analysis was done as previously reported (39, 40). Overnight cultures were diluted to OD_600_ 0.05 in fresh LB or TSB. For growth measurements, 100 µL aliquots were dispensed into 96-well plates and incubated at 37 °C, and OD_600_ was recorded hourly for 14 h. For morphology, cultures grown in LB, TSB, or LB_0_N were diluted to OD_600_ 0.05 and grown to mid-log phase (OD_600_ 0.4–0.5). Cells were labeled with HADA to visualize sites of PG synthesis, fixed with paraformaldehyde, and mounted on 1.5% agarose pads in 1× PBS. Imaging was performed on a Nikon Eclipse Ti-2 wide-field epifluorescence microscope equipped with a Photometrics Prime 95B camera and a Plan Apo 100×/1.45 NA objective. Images were acquired with NIS-Elements and analyzed in ImageJ.

### Determination of MICs

MICs were measured using E-test strips (Liofilchem) as described (12, 22). Briefly, strains were spread uniformly on LB agar plates using sterile swabs. After the surface dried, an E-test strip was applied, plates were incubated at 37 °C for 18–20 h, and MICs were read at the intersection of the inhibition ellipse with the strip scale.

### Isolation of LOS⁻ *A. baumannii* and determination of mutation frequency

LOS⁻ isolation was performed as described (12, 13). Cultures were grown in LB at 37 °C to mid-log or stationary phase. Where indicated, complementation strains were grown with kanamycin (25 µg/mL) or hygromycin (250 µg/mL) and IPTG (1 mM). One milliliter of culture at OD_600_ 1.0 (approximately 10⁹ CFU) was harvested (8,000 rpm, 2 min), washed once in LB, and plated on LB agar containing colistin (10 µg/mL). For complementation strains, plates also contained the corresponding antibiotic and IPTG. Colonies were replica-plated onto LB agar containing vancomycin (10 µg/mL) or colistin (10 µg/mL). Colonies that were colistin resistant and vancomycin sensitive were classified as LOS⁻. Mutation frequency was calculated as the number of LOS⁻ colonies divided by the total viable CFUs plated.

### RNA sequencing

RNA-seq was performed as described (29, 40, 45) with minor modifications. Total RNA was extracted from cultures grown in LB to OD_600_ 0.4–0.5 (three biological replicates per condition) using the Direct-Zol RNA Miniprep Kit (Zymo Research). Genomic DNA was removed using TURBO DNA-free (Invitrogen). Libraries were sequenced at SeqCenter on an Illumina NextSeq 550 platform. Reads were mapped to the ATCC 17978 reference genome using CLC Genomics Workbench (Qiagen), and expression values were calculated as RPKM. Differential expression analyses were performed across strains, and plots were generated in GraphPad Prism. Sequencing data were deposited in GEO under accession number GSE314574.

### Western blotting

Western blotting was performed as described (12, 39, 45). Proteins were separated by SDS-PAGE and transferred to PVDF membranes (Thermo Fisher Scientific). Membranes were blocked for 2 h at room temperature in Tris-buffered saline containing 5% nonfat milk. Primary antibodies were incubated overnight at 4 °C (α-PBP1A, 1:1,000; α-RpoA, 1:1,000), followed by HRP-conjugated α-rabbit secondary antibody (1:10,000; Thermo Fisher Scientific). Signal was developed using Super Signal West Pico Plus (Thermo Fisher Scientific) and imaged under no saturating conditions to enable comparison of relative protein abundance.

### Quantification and Statistical Analysis

Unless otherwise indicated, all experiments were performed using at least three independent biological replicates, defined as cultures grown and processed on separate days from independent starter cultures. Data are presented as mean ± standard deviation (SD).

Quantification of colony-forming units (CFUs), LOS-deficient variant frequencies, and phospholipid abundances was performed using ImageJ or manual enumeration, as indicated in the figure legends. For radiolabeled phospholipid analyses, band intensities from thin-layer chromatography (TLC) plates were quantified in ImageJ and expressed as a percentage of total signal per lane. Immunoblot band intensities were compared qualitatively under nonsaturating exposure conditions and were not subjected to formal statistical testing.

RNA-seq data were generated from three biological replicates per condition. Differential gene expression analysis was performed using CLC Genomics Workbench (Qiagen). Genes were considered significantly differentially expressed if they met the criteria of log₂ fold change ≥ 1 and false discovery rate (FDR) < 0.05. Volcano plots and heat maps were generated in GraphPad Prism.

Statistical comparisons between two groups were performed using unpaired two-tailed Student’s *t*-tests when data followed an approximately normal distribution. Experiments were not randomized or blinded, and no statistical methods were used to predetermine sample size. Exact values of *n*, the definition of replicates, statistical tests used, and significance thresholds are specified in the relevant figure legends.

## Acknowledgments

The work was supported by funding from the National Institutes of Health (grants R35GM143053 and R01AI168159 to J.M.B). The funders had no role in study design, data collection and analysis, decision to publish, or preparation of the manuscript.

## Supplementary Tables and Figure Legends

**Table S1:** Differential gene expression profiles in outer membrane homeostasis mutants associated with colistin resistance.

**Table S2:** Strains and plasmids used in this study.

**Table S3:** Primers used in this study.

---

**Figure S1.**
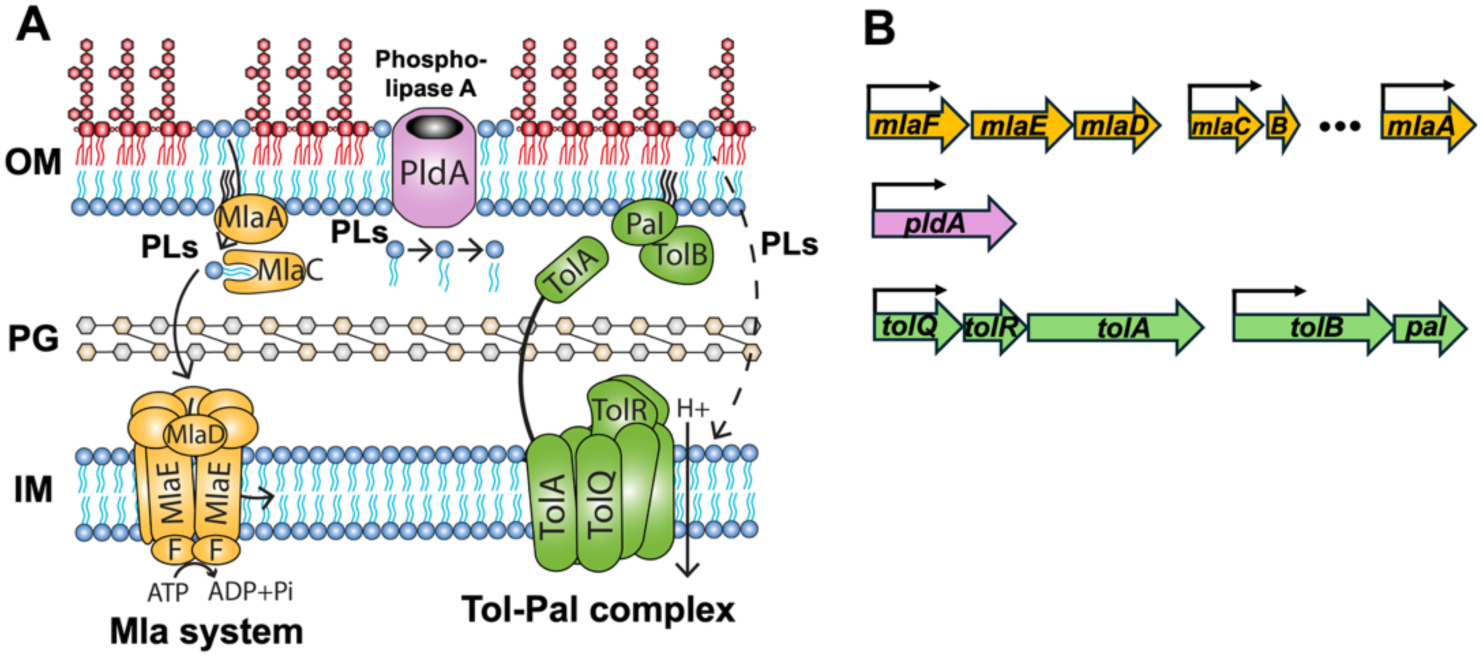
Systems that maintain OM lipid homeostasis in *A. baumannii*. (A) Diagram showing the three main systems that contribute to OM lipid homeostasis in *A. baumannii*. These include the Mla system (orange), which mediates retrograde PLs transport; the OM phospholipase PldA (purple), which degrades mislocalized PLs; and the Tol-Pal complex (green), which ensures proper OM and PG connection during cell division. (B) Genetic arrangement of each system. The *mlaFEDCB* operon (genes *A1S_3103–3099*), encoding the core components of the Mla transport machinery, was deleted entirely. The gene *mlaA* (A1S_0622), which encodes an OM-anchored lipoprotein, is located elsewhere in the chromosome and was not disrupted in this study. *pldA* is a monocistronic gene (*A1S_1919*). The *tol-pal* system is organized into two adjacent operons *tolQRA* (*A1S_2591–2593*), and *tolB-pal* (*A1S_2594–2595*), which were deleted together as a functional unit. This organization enabled the systematic construction of all single, double, and triple mutant combinations.

**Figure S2.**
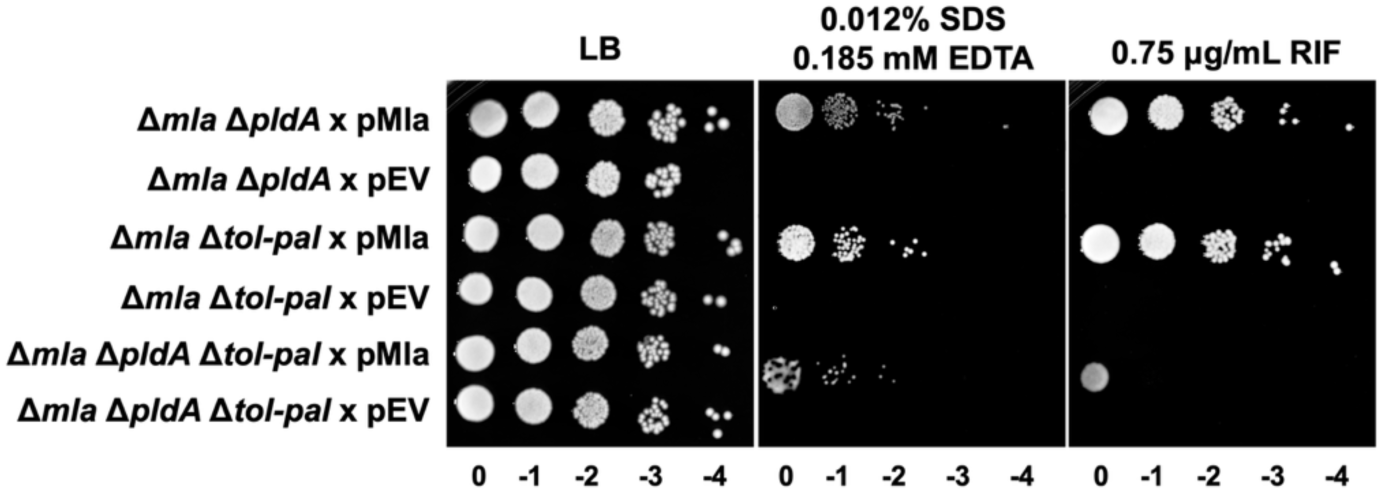
Expression of Mla partially restores envelope integrity in double and triple mutants. Serial dilution assay on LB agar containing SDS and EDTA demonstrates that expression of the *mla* operon partially rescues detergent resistance in Δ*mla* Δ*pldA*, Δ*mla* Δ*tol*-*pal*, and Δ*mla* Δ*pldA* Δ*tol*-*pal* mutant strains. Rifampicin susceptibility assay shows partial restoration of resistance upon *mla* expression in the same mutant backgrounds.

**Figure S3.**
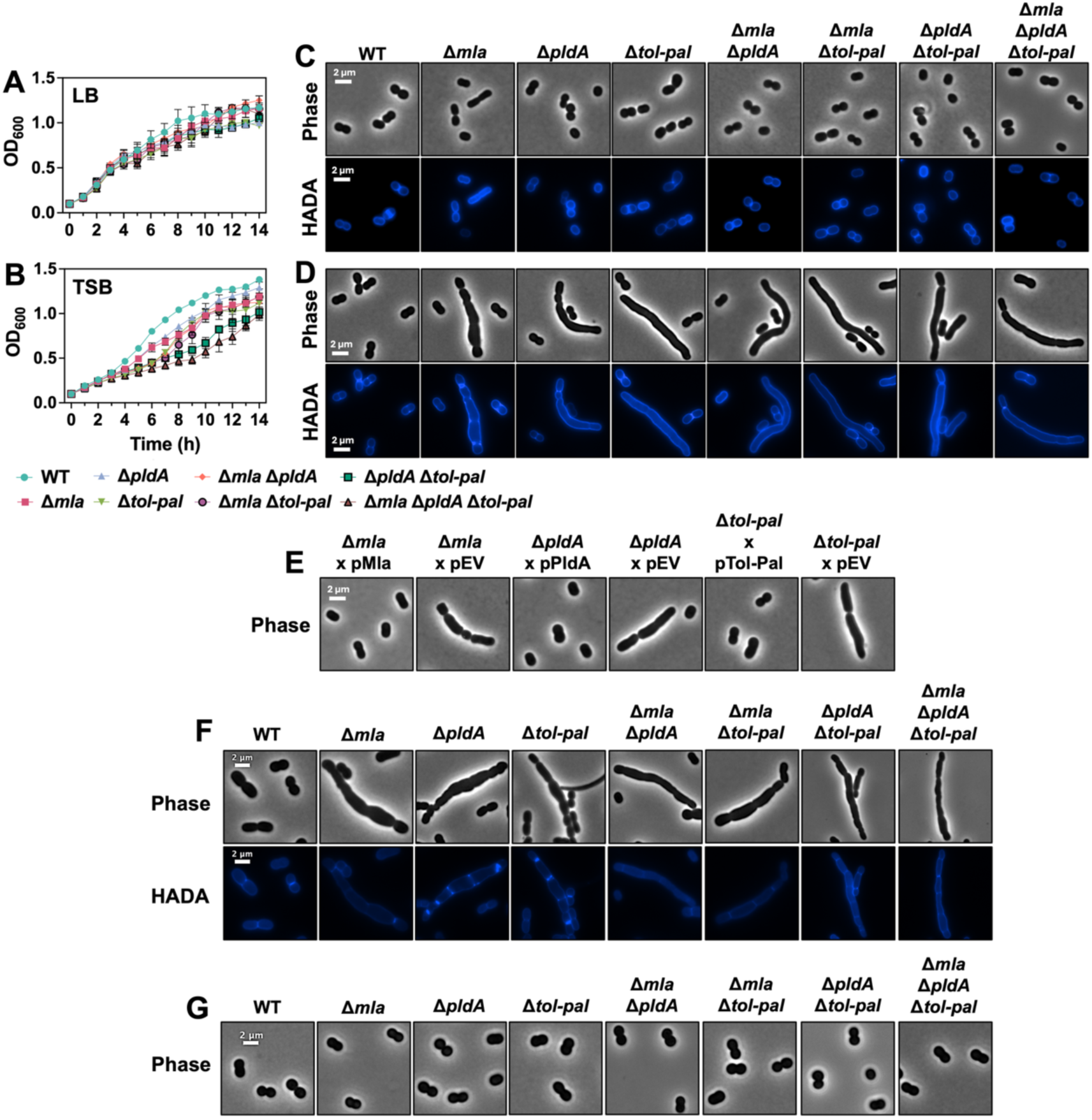
Disruption of OM lipid homeostasis impairs growth and cell division under envelope stress conditions. (A-B) Growth curves for wild-type (*A. baumannii* ATCC 17978, WT) and mutant strains in LB (A) and TSB (B). Cultures were inoculated at OD_600_ 0.05 and monitored over time. (C-D) Phase-contrast and HADA fluorescence microscopy of WT and mutant strains grown in LB (C) or TSB (D). Mutants exhibited cell elongation and septation defects in TSB. Scale bars: 2 µm. Growth curves represent three independent experiments; microscopy images correspond to one representative experiment. (E) Phase-contrast microscopy of cells grown in TSB medium shows that in *trans* expression of *mla*, *pldA*, or *tol*-*pal* partially rescues the morphological defects observed in the corresponding deletion mutants. Complemented strains regain near-normal rod shape, while empty vector controls exhibit marked cell elongation and division defects. (F) Phase-contrast and HADA fluorescence microscopy of wild-type (WT) and mutant strains grown in LB_0_N. (G) Growth in LB_0_N supplemented with 171 mM sucrose reproduces similar morphological stabilization. These results support the conclusion that osmotic imbalance contributes to envelope instability in OM homeostasis mutants. WT cells maintain a uniform rod-shaped morphology under all conditions.

**Figure S4.**
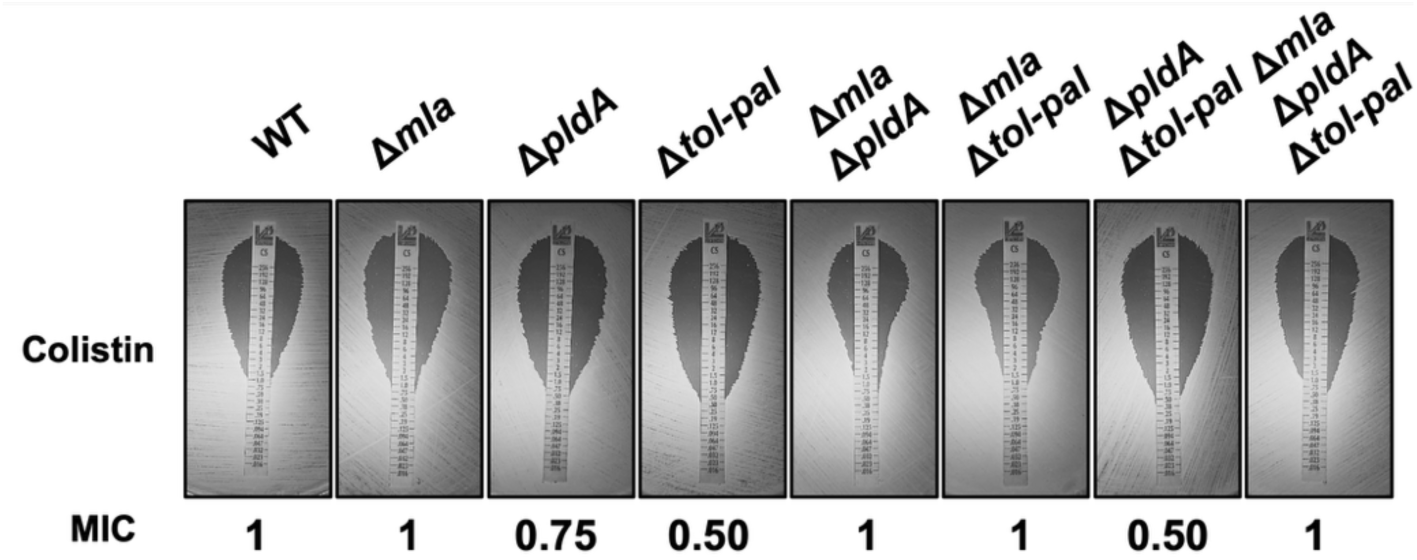
Colistin susceptibility of *A. baumannii* mutants determined by E-test. Minimum inhibitory concentrations (MICs) for colistin were determined using E-test strips on LB agar plates for wild-type (WT) and mutant strains. WT, Δ*mla*, Δ*mla* Δ*pldA*, Δ*mla* Δ*tol*-*pal*, and the Δ*mla* Δ*pldA* Δ*tol*-*pal* triple mutant display comparable MIC values. In contrast, Δ*pldA* and Δ*tol*-*pal* single and double mutants exhibited moderately reduced MICs, suggesting increased susceptibility to colistin in the absence of individual components involved in lipid homeostasis or envelope stability components.

**Figure S5.**
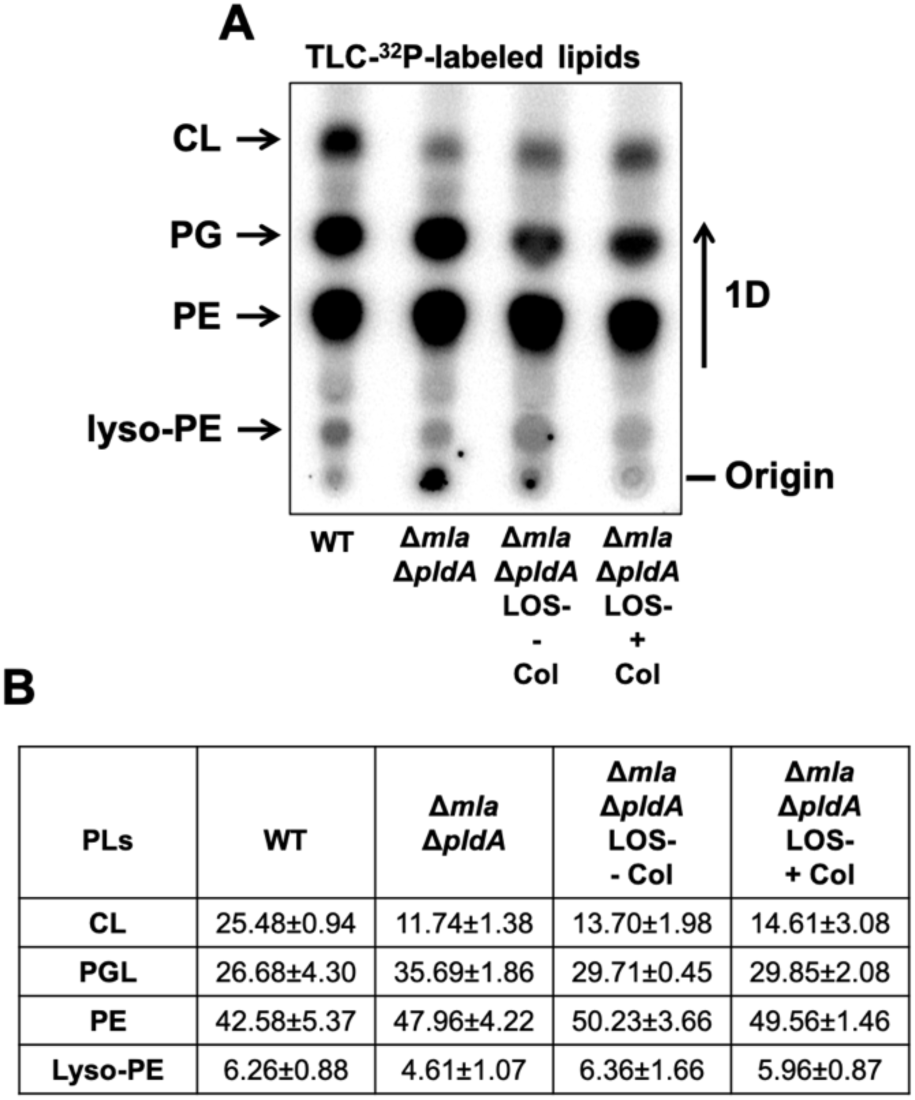
PL composition of colistin-selected LOS-deficient derivatives. (A) Thin-layer chromatography (TLC) analysis of ^^32^P-labeled PLs extracted from wild-type (WT), Δ*mla* Δ*pldA*, and Δ*mla* Δ*pldA* LOS-deficient derivatives propagated in the absence (− colistin) or presence (+ colistin) of colistin. Lipid species were resolved by one-dimensional TLC and visualized by phosphorimaging. Positions of cardiolipin (CL), phosphatidylglycerol (PG), phosphatidylethanolamine (PE), and lyso-PE are indicated. (B) Quantification of major PL species expressed as percentage of total signal. Data represent the mean of three independent experiments ± standard deviation.

**Figure S6.**
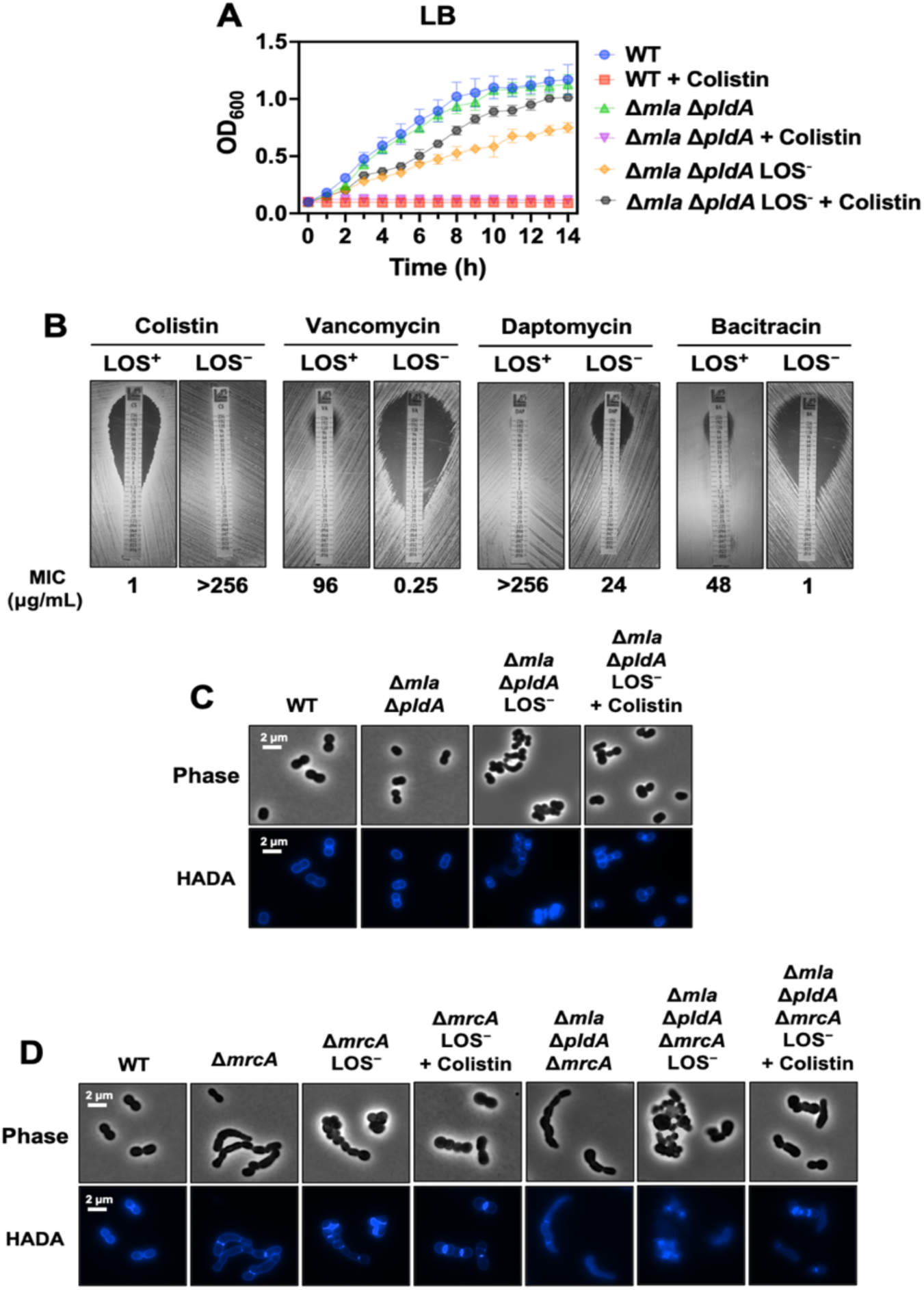
Morphological and growth characterization of colistin-selected LOS-deficient variants under colistin exposure. (A) Growth curves of wild-type (WT), the Δ*mla* Δ*pldA* parental mutant, and colistin-selected LOS-deficient variant in LB medium with or without 10 μg/mL colistin. The LOS-deficient variant exhibits impaired growth in the absence of antibiotic but shows enhanced growth upon colistin treatment, which is lethal to both WT and its parental strain. (B) Antibiotic susceptibility profiling using E-test strips shows that the Δ*mla* Δ*pldA* parental strain remains susceptible to colistin but resistant to large hydrophilic antibiotics such as vancomycin, daptomycin, and bacitracin. Values indicate (μg/mL). In contrast, the LOS-deficient variant exhibits high-level colistin resistance along with increased susceptibility to these antibiotics, reflecting lipid A loss and compromised OM integrity. (C) Phase-contrast and HADA fluorescence microscopy of WT, Δ*mla* Δ*pldA*, the LOS-deficient variant, and the LOS-deficient variant treated with colistin. WT and Δmla Δ*pldA* cells display typical coccobacillary morphology with defined septa. In contrast, the LOS-deficient variant shows marked elongation, curvature, septation defects, and extensive cellular aggregation. Colistin exposure partially restores normal morphology, reduces aggregation, and improves septal organization. Scale bars: 2 µm. Images are representative of three independent experiments. (D) Phase-contrast and HADA fluorescence microscopy of Δ*mrcA*, Δ*mrcA* LOS-deficient derivatives, and Δ*mla* Δ*pldA* Δ*mrcA* LOS-deficient variants, with and without colistin treatment. LOS-deficient cells arising from both genetic backgrounds exhibit pronounced morphological defects, including elongation and septation abnormalities, consistent with those observed in other LOS-deficient states. Images are representative of three independent experiments.

