## Supplemental Material for "Outer membrane remodeling via lipid-peptidoglycan crosstalk enables lipooligosaccharide-deficient colistin resistance"

**Supplementary Figures**
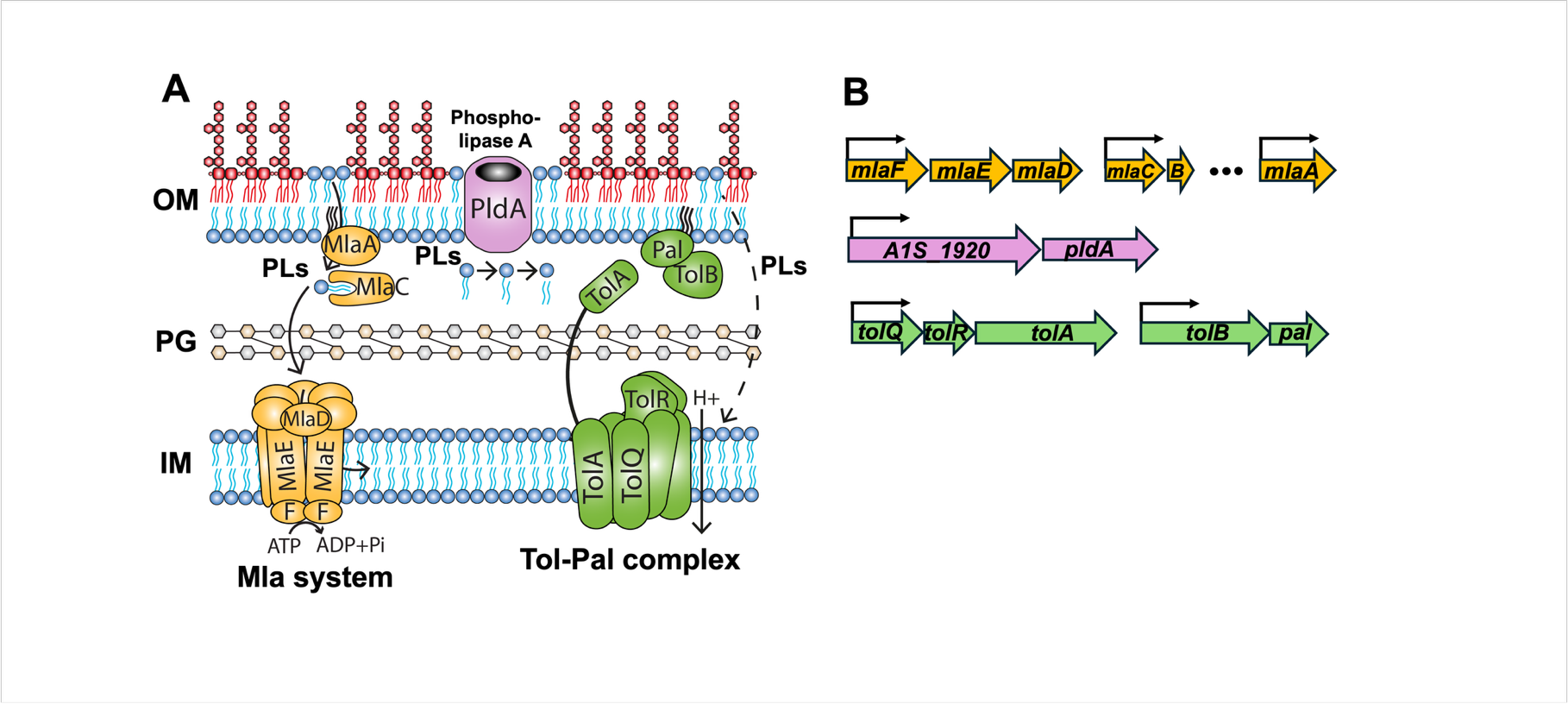

**Figure S1. Systems that maintain OM lipid homeostasis in *A. baumannii*.** (A) Diagram showing the three main systems that contribute to OM lipid homeostasis in *A. baumannii*. These include the Mla system (orange), which mediates retrograde PLs transport; the OM phospholipase PldA (purple), which degrades mislocalized PLs; and the Tol-Pal complex (green), which ensures proper OM and peptidoglycan connection during cell division. (B) Genetic arrangement of each system. The *mlaFEDCB* operon (genes *A1S_3103–3099*), encoding the core components of the Mla transport machinery, was deleted entirely. The gene *mlaA* (*A1S_0622*), which encodes an OM-anchored lipoprotein, is located elsewhere in the chromosome and was not disrupted in this study. *pldA* (*A1S_1919*) is the second gene in a predicted operon with *A1S_1920*. The *tol-pal* system is organized into two adjacent operons *tolQRA* (*A1S_2591*–*2593*), and *tolB-pal* (*A1S_2594*–*2595*), which were deleted together as a functional unit. This organization enabled the systematic construction of all single, double, and triple mutant combinations.

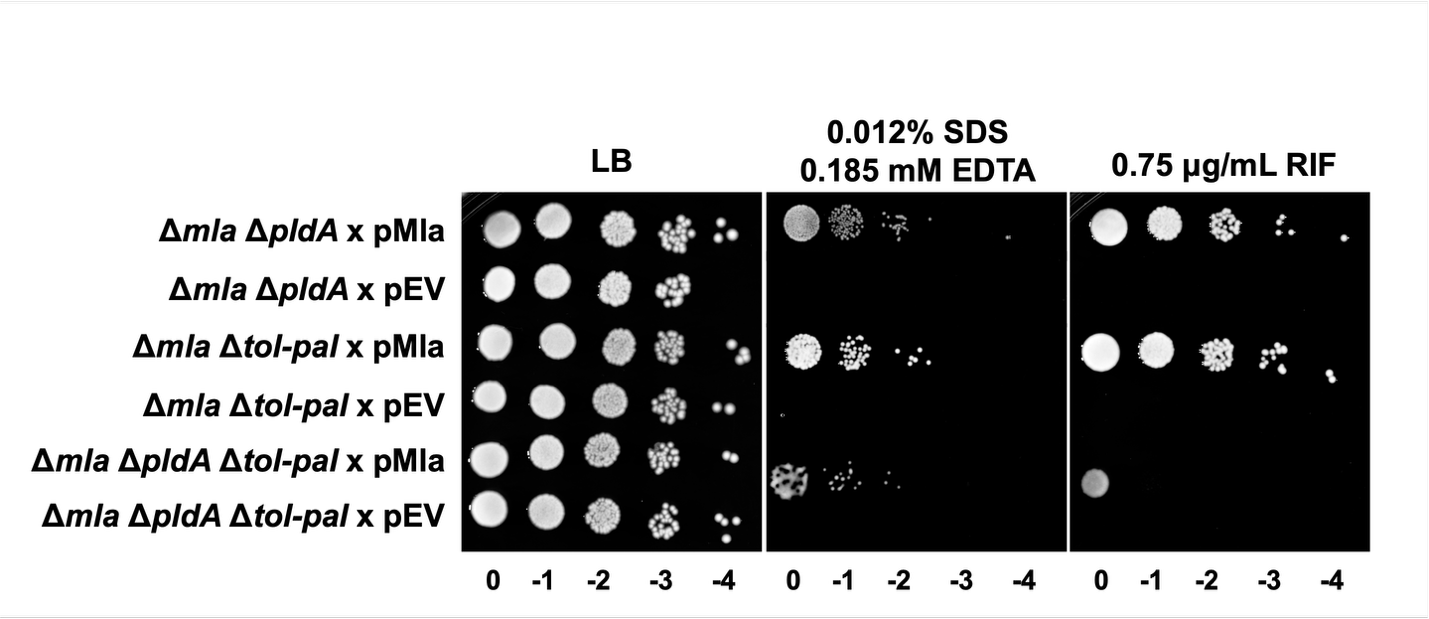

**Figure S2. Expression of Mla partially restores envelope integrity in double and triple mutants.** Serial dilution assay on LB agar containing SDS and EDTA demonstrates that expression of the *mla* operon partially rescues detergent resistance in Δ*mla* Δ*pldA*, Δ*mla* Δ*tol*-*pal*, and Δ*mla* Δ*pldA* Δ*tol*-*pal* mutant strains. Rifampicin (RIF) susceptibility assay shows partial restoration of resistance upon *mla* expression in the same mutant backgrounds. Data shown are representative of three independent experiments.

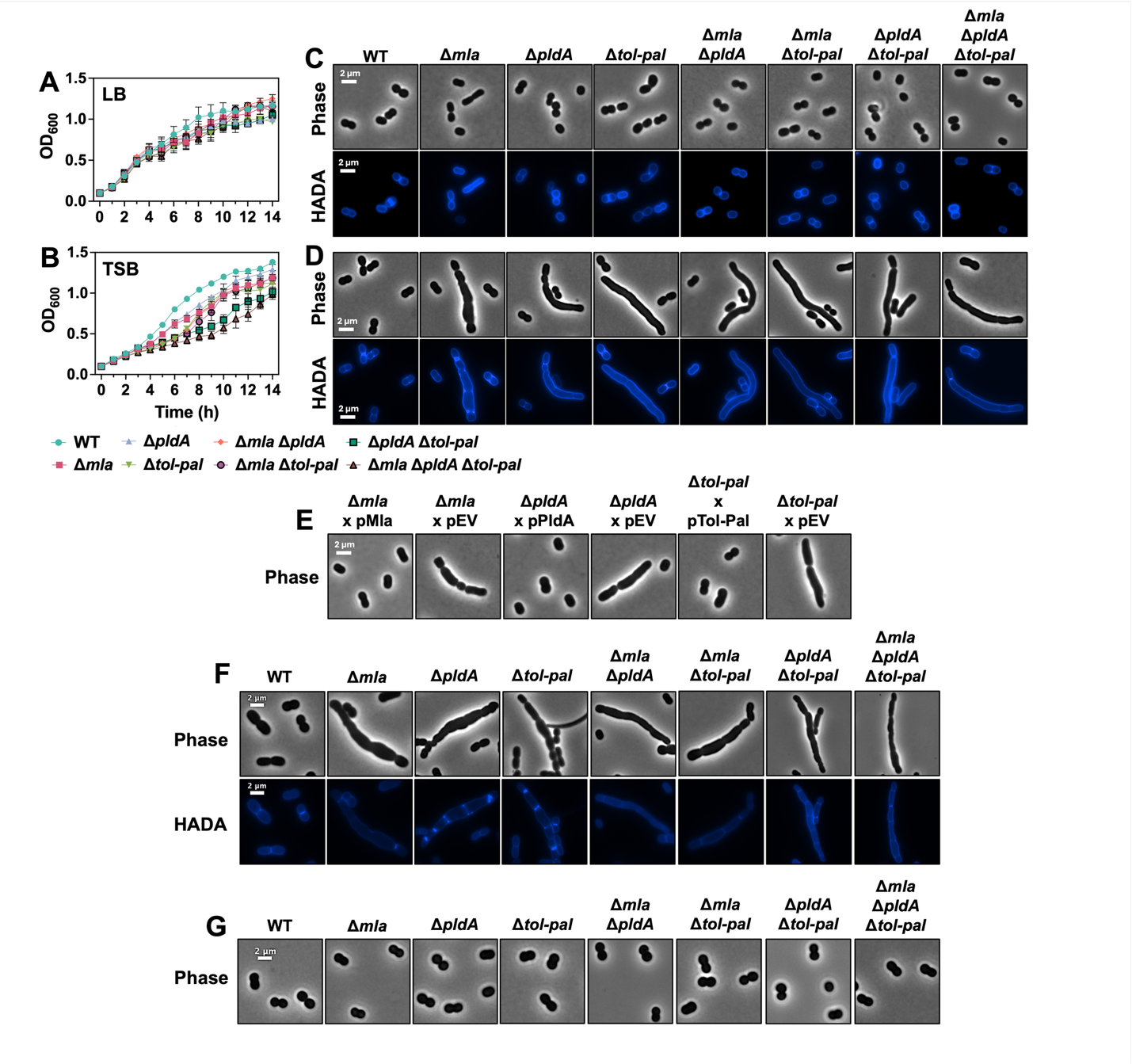

**Figure S3. Disruption of OM lipid homeostasis impairs growth and cell division under envelope stress conditions.** (A-B) Growth curves for wild-type (*A. baumannii* strain ATCC 17978, WT) and mutant strains in LB (A) and TSB (B). Cultures were inoculated at OD_600_ 0.05 and monitored over time. (C-D) Phase-contrast and HADA fluorescence microscopy of WT and mutant strains grown in LB (C) or TSB (D). Mutants exhibited cell elongation and septation defects in TSB. Scale bars: 2 µm. Growth curves represent three independent experiments; microscopy images correspond to one representative experiment. (E) Phase-contrast microscopy of cells grown in TSB medium shows that in *trans* expression of *mla*, *pldA*, or *tol*-*pal* partially rescues the morphological defects observed in the corresponding deletion mutants. Scale bars: 2 µm. (F) Phase-contrast and HADA fluorescence microscopy of wild-type (WT) and mutant strains grown in LB_0_N. (G) Growth in LB_0_N supplemented with 171 mM sucrose reproduces WT morphology. These results support the conclusion that osmotic imbalance contributes to envelope instability in OM homeostasis mutants. WT cells maintain a uniform rod-shaped morphology under all conditions.

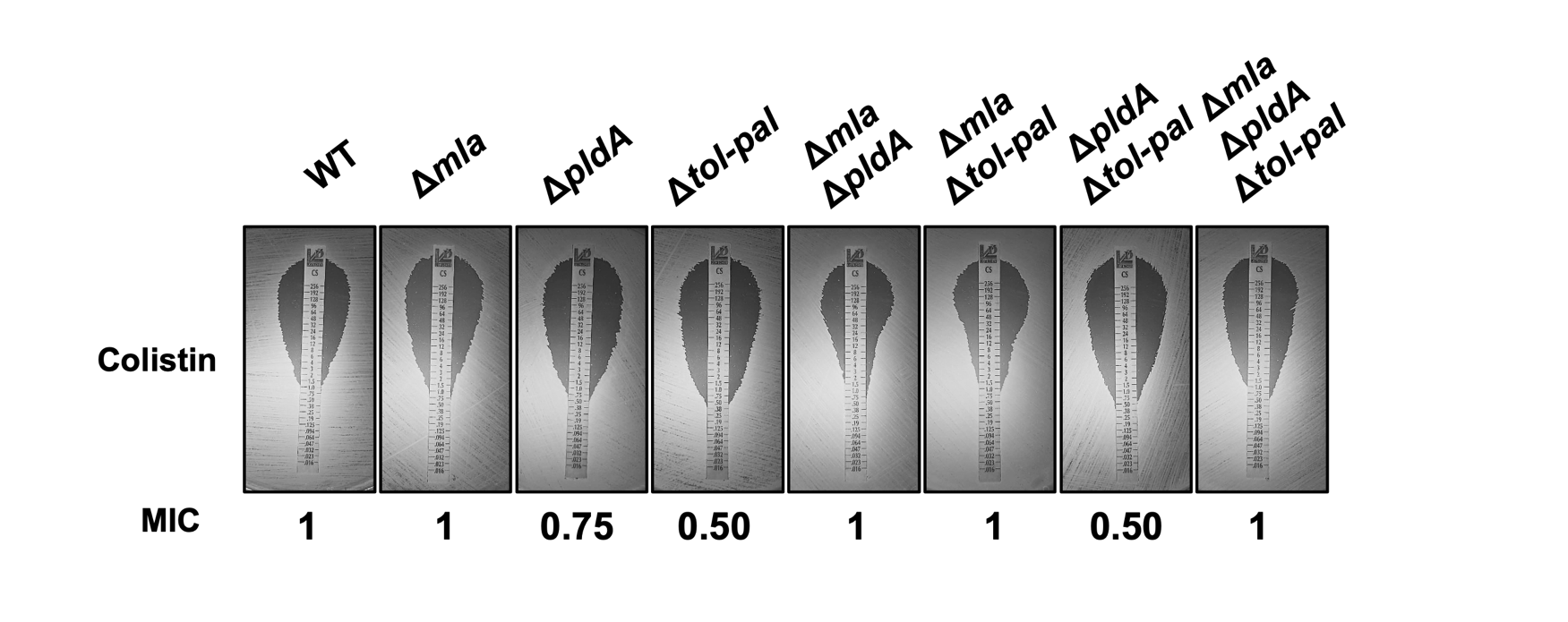

**Figure S4. Colistin susceptibility of *A. baumannii* mutants determined by E-test.** Minimum inhibitory concentrations (MICs) for colistin were determined using E-test strips on LB agar plates for wild-type (WT) and mutant strains. WT, Δ*mla*, Δ*mla* Δ*pldA*, Δ*mla* Δ*tol*-*pal*, and the Δ*mla* Δ*pldA* Δ*tol*-*pal* triple mutant display comparable MIC values. In contrast, Δ*pldA* and Δ*tol*-*pal* single and double mutants exhibited moderately reduced MICs, suggesting slightly increased susceptibility to colistin in the absence of individual components involved in lipid homeostasis or envelope stability components.

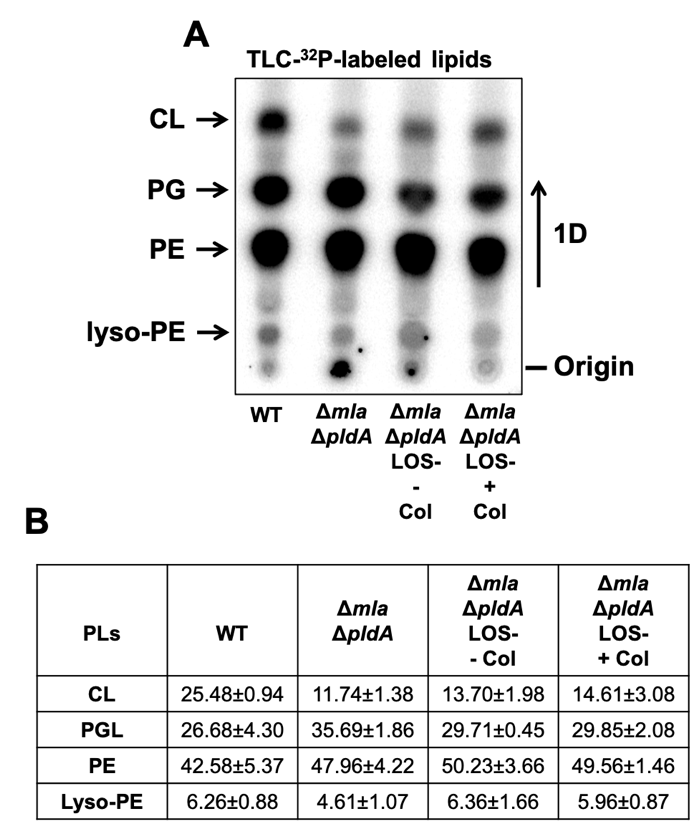

**Figure S5. PL composition of colistin-selected LOS-deficient derivatives.** (A) Thin-layer chromatography (TLC) analysis of ^^32^P-labeled PLs extracted from wild-type (WT), Δ*mla* Δ*pldA*, and Δ*mla* Δ*pldA* LOS-deficient derivatives propagated in the absence (− colistin) or presence (+ colistin) of colistin. Lipid species were resolved by one-dimensional TLC and visualized by phosphorimaging. Positions of cardiolipin (CL), phosphatidylglycerol (PG), phosphatidylethanolamine (PE), and lyso-PE are indicated. (B) Quantification of major PL species expressed as percentage of total signal. Data represent the mean of three independent experiments ± standard deviation.

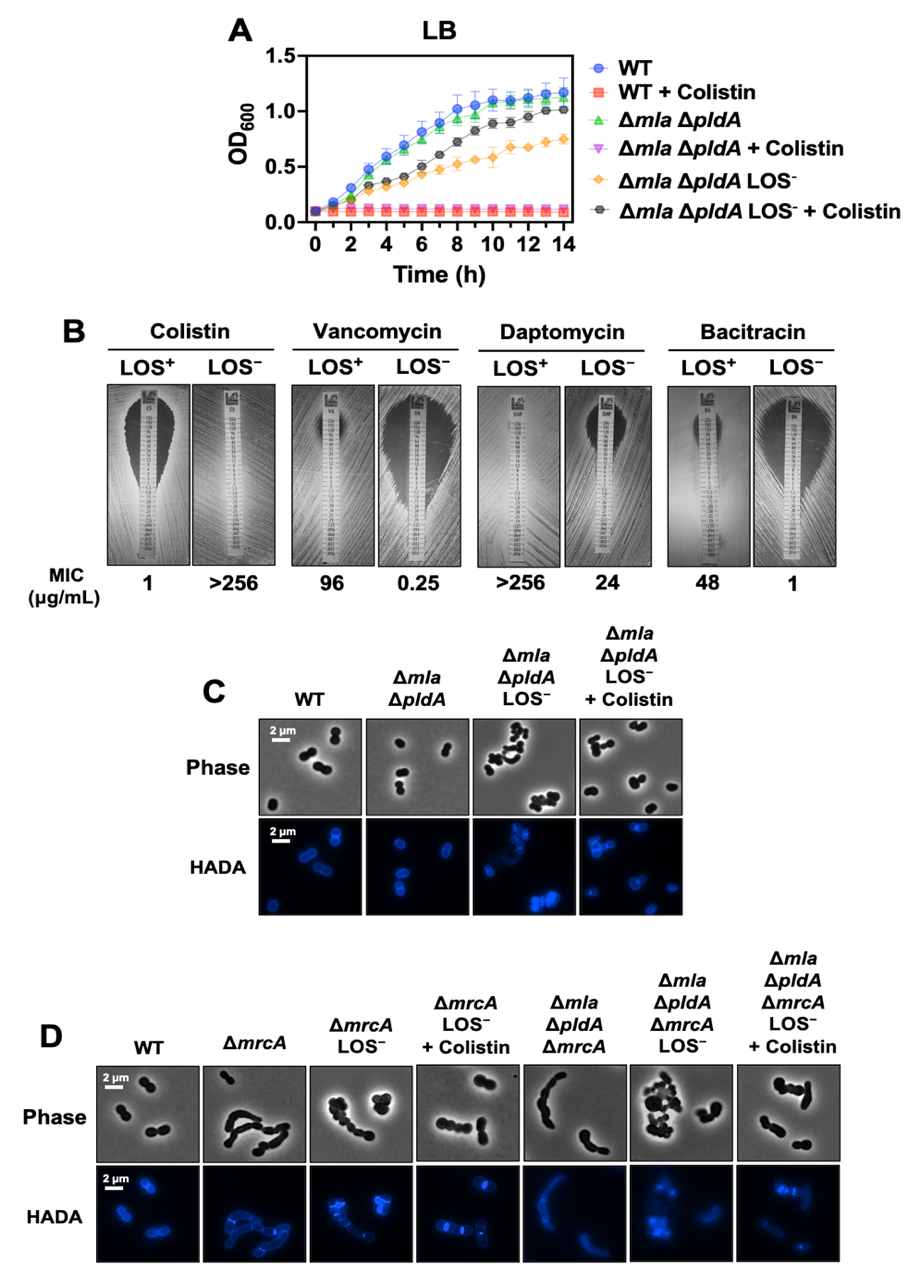

**Figure S6. Morphological and growth characterization of colistin-selected LOS-deficient variants.** (A) Growth curves of wild-type (WT), the Δ*mla* Δ*pldA* parental mutant, and colistin-selected LOS-deficient variant in LB medium with or without 10 μg/mL colistin. LOS-deficient variant exhibits impaired growth in the absence of antibiotic but shows enhanced growth upon colistin treatment, which is lethal to both WT and its parental strain. (B) Antibiotic susceptibility profiling using E-test strips shows that the Δ*mla* Δ*pldA* parental strain remains susceptible to colistin but resistant to large hydrophilic antibiotics such as vancomycin, daptomycin, and bacitracin. Values indicate (μg/mL). In contrast, the LOS-deficient variant exhibits high-level colistin resistance along with increased susceptibility to these antibiotics, reflecting lipid A loss and compromised OM integrity. (C) Phase-contrast and HADA fluorescence microscopy of WT, Δ*mla* Δ*pldA*, the LOS-deficient variant, and the LOS-deficient variant treated with colistin. WT and Δ*mla* Δ*pldA* cells display typical coccobacillary morphology with defined septa. In contrast, the LOS-deficient variant shows marked elongation, curvature, septation defects, and extensive cellular aggregation. Colistin exposure partially restores normal morphology, reduces aggregation, and improves septal organization. Scale bars: 2 µm. Images are representative of three independent experiments. (D) Phase-contrast and HADA fluorescence microscopy of Δ*mrcA*, Δ*mrcA* LOS-deficient derivatives, and Δ*mla* Δ*pldA* Δ*mrcA* LOS-deficient variants, with and without colistin treatment. LOS-deficient cells arising from both genetic backgrounds exhibit pronounced morphological defects, including elongation and septation abnormalities, consistent with those observed in other LOS-deficient states. Images are representative of three independent experiments.

**Supplementary Tables**

**Please see the excel spreadsheet (separate file) for Table S1.**

**Table S2: Strains and plasmids used in this study.**

| **Strain or Plasmid** | **Description** | **Reference or Source** |
| --- | --- | --- |
| **Strains** |  |  |
| *A. baumannii* ATCC 17978 | Wild type | (1) |
| *A. baumannii* ATCC 17978 | *Δmla* | This study |
| *A. baumannii* ATCC 17978 | *ΔpldA* | This study |
| *A. baumannii* ATCC 17978 | *Δtol-pal* | This study |
| *A. baumannii* ATCC 17978 | *Δmla ΔpldA* | This study |
| *A. baumannii* ATCC 17978 | *Δmla Δtol-pal* | This study |
| *A. baumannii* ATCC 17978 | *ΔpldA Δtol-pal* | This study |
| *A. baumannii* ATCC 17978 | *Δmla ΔpldA Δtol-pal* | This study |
| *A. baumannii* ATCC 17978 | *ΔmrcA* | This study |
| *A. baumannii* ATCC 17978 | *Δmla ΔpldA ΔmrcA* | This study |
| **Plasmids** |  |  |
| pAT03 | pMMB67EH with FLP recombinase, Tet^R^ | (2) |
| pAT04 | pMMB67EH with REC*_Ab_* system, Tet^R^ | (2) |
| pKD4 | Km^R^ | (3) |
| pMMB67EH-Km | pMMB67EH with the Km^R^ gene from pKD4 inserted into the PvuI site, Km^R^ | (4) |
| pABBR-Km | pABBR-Km-*mrcA* (Native promoter) | (4) |
| pJMP3665-Hyg | Hyg^R^ | (5) |
| pMMB67EH-Km | pMMB67EH-Km-*mla* (ITPG) | This study |
| pMMB67EH-Km | pJMP3665-Hyg-*pldA* (IPTG) | This study |
| pMMB67EH-Km | pMMB67EH-Km-*tol-pal* (ITPG) | This study |
| pMMB67EH-Km | pMMB67EH-Km-*mrcA*-Chis (IPTG) | This study |

**Table S3: Primers used in this study.**

| **Oligo Name** | **Sequence (5’ to 3’)** |
| --- | --- |
| **Deletion Primers** |  |
| 17978 *mlaFEDCB* Kan FRT 5' | ATGAATAATAAAACTCCTCTCTCGACTCAATCACTGATTGAAGTGAAAAACTTGAGTTTTAACCGGGGGAACGCGTCATTTATGACAATATTAGTTTAAATATACGTCGAGGCCAGATCACGGCAGCGATTGTGTAGGCTGGAGCTGCTTCG |
| 17978 *mlaFEDCB* Kan FRT 3' | TCAAACCAAATGTAGGTCTTCTTGCAAATGGCAGGCTTG AATGATTTTCAGCATCTTCTCCGGTACATTTTTAAAATGTAACCCAGACTT TTGTGGAGTTTGACGTAGCCATTGCAC TAAAACAGATATCCTCCTTAGTTCCTATTCCG |
| 17978 *mlaFEDCB* confirm 5' | GAGTCGCCTTTGGCATGGTTG |
| 17978 *mlaFEDCB* confirm 3' | CTCGCTTGTCGAACATGACGC |
| 17978 *pldA* Kan FRT 5' | ATGCCGACACCTTAGCCCCTGTAAAACCTGCGAGTGTAGATGCATGTGTTGCCCTTGCTTCGAATGCTGATCGCCTAGCTTGTTATGATGCTGTGTTTAAACCATCGGCCTTACCAGTAGTTCAAAGCGATTGTGTAGGCTGGAGCTGCTTCG |
| 17978 *pldA* Kan FRT 3' | TAAGTTGCACGATGGTTATAGTCAATTAAGCTTTCACCATAACCATTAAATAATTGGAAGTGTCCGCGCAGCTTGCCACTAATCGGGAAAGCCCAGTCAAACTGAACAGCACCATGAGAATCATCATATCCTCCTTAGTTCCATTTCCG |
| 17978 *pldA* 5' confirm | CACCAGCAGGTGAAGCAATAACAG |
| 17978 *pldA* 3' confirm | CCTGTACAGTAGGGCTTTCGCG |
| 17978 *tol-pal* Kan FRT 5' | ATGGCAACAAACATTGAATCAACCCTGCATATTTCTGACCTTATTTTACAAGCAAGTCCAGTCGTCCAGTTGGTCATGCTGATTTTATTGTTAGCATCAATTTTTAGTTGGTACCTGATTGCCAAAGCGATTGTGTAGGCTGGAGCTGCTTCG |
| 17978 *tol-pal* Kan FRT 3' | TTATTTTAATAGAGGAGGAACCGCTTCATAGTTAATTTCAACGCGGCGGTTTTCTTTCCAAGCTGATTCATCATGGCCAGGATTAACAGGCGCTTCTTTACCATAACTTACAGCTTCAAGTTGCTATATCCTCCTTAGTTCCTATTCCG |
| 17978 *tol-pal* 5' confirm | CGCTATATGGAACGTACCCGTACAG |
| 17978 *tol-pal* 3' confirm | GACTTGAGCCAACTCAGGTGTAAG |
| 17978 *mrcA* Kan FRT 5' | ATGAAAAAGCTATCCAGTTTGGGCTTCGTGCGTCCAATTTTTTTGATCATTATTATTATCTTAGTCTCACTTCCGATGGGATTCTATGGCATGTATCTCTATATTGCCCCAACGCTTCCGGAAATAGCGATTGTGTAGGCTGGAGCTGCTTCG |
| 17978 *mrcA* Kan FRT 3' | TTACTCAATTTGATTAATCAAGTTTTCTAATTCATCTTTTTCACGCTTCGGCGCTGCCCCTTGCGAAGGTTTCATCGCTGGAGGTGTTGTCTTGGAAGGAATCACAATTTCCTCACCAGGTAAGTATATCCTCCTTAGTTCCTATTCCG |
| 17978 *mrcA* 5' confirm | CGGGTAAAGGCATCAAAGCATAGC |
| 17978 *mrcA* 3' confirm | CAGGAGCAACTTTACCTGTCATCG |
| **Complementation Primers** |  |
| 17978 *mlaFEDCB* KpnI 5' | CGCGGTACCATGAATAATAAAACTCCTCTCTCGACTC |
| 17978 *mlaFEDCB* SalI 3' | CGCGTCGACTCAAACCAAATGTAGGTCTTCTTGC |
| 17978 *pldA* BamHI 5' | CGCGGATCCATGGCGTTCAGACAATTTGAACGC |
| 17978 *pldA* XbaI 3' | CGCTCTAGATTAGTACCAGTTCATGAGTGAAACACCCAAG |
| 17978 *tol-pal* BamHI 5' | CGCGGATCCATGGCAACAAACATTGAATCAACCCTG |
| 17978 *tol-pal* SalI 3' | CGCGTCGACTTATTTTAATAGAGGAGGAACCGCTTCATAG |
| 17978 *mrcA* KpnI 5' | CGCGGTACCATGAAAAAGCTATCCAGTTTGGGCTTCGTGC |
| 17978 *mrcA* 6xHis BamHI 3' | CGCGGATCCTTAATGGTGATGGTGATGGTGTTCAATTTGATTAATCAAGTTTTCTAATTCATCTTTTTCAC |
